# Dichotomous Feedback: A Signal Sequestration-based Feedback Mechanism for Biocontroller Design

**DOI:** 10.1101/2021.12.27.474252

**Authors:** Aivar Sootla, Nicolas Delalez, Emmanouil Alexis, Arthur Norman, Harrison Steel, George H. Wadhams, Antonis Papachristodoulou

## Abstract

We introduce a new design framework for implementing negative feedback regulation in Synthetic Biology, which we term ‘dichotomous feedback’. Our approach is different from current methods, in that it sequesters existing fluxes in the process to be controlled, and in this way takes advantage of the process’s architecture to design the control law. This signal sequestration mechanism appears in many natural biological systems and can potentially be easier to realise than ‘molecular sequestration’ and other comparison motifs that are nowadays common in biomolecular feedback control design. The loop is closed by linking the strength of signal sequestration to the process output. Our feedback regulation mechanism is motivated by two-component signalling systems, where we introduce a second response regulator competing with the natural response regulator thus sequestering kinase activity. Here, dichotomous feedback is established by increasing the concentration of the second response regulator as the level of the output of the natural process increases. Extensive analysis demonstrates how this type of feedback shapes the signal response, attenuates intrinsic noise while increasing robustness and reducing crosstalk.

---

Synthetic Biology has made many recent advances, including the engineering of genetically modified organisms to produce new compounds, detect harmful substances or for drug delivery [Brophy and Voigt, 2014, Purnick and Weiss, 2009, Freemont and Kitney, 2012, Din et al., 2016, Ozdemir et al.,2018]. While these and future applications offer great promise, there are a number of challenges that Synthetic Biology faces. One of them is the reliable and systematic design of synthetic circuits which is made difficult by noise, burden, and crosstalk inherent to biological systems.

Feedback control is often used for modulating the response and improving the robustness of synthetic biological circuits [Del Vecchio and Murray, 2014, Hsiao et al., 2018], in much the same way negative feedback is used in natural biological and technological systems [Thomas and d’Ari, 1990] so that they can respond effectively to noise and disturbances in their environment. Consider, for example, how the altitude of an aircraft is kept constant despite wind and other disturbances [Astrom and Murray, 2010]; or how the bacterial chemotaxis system [Yi et al., 2000] or the heat shock feedback control system [El-Samad et al., 2005] operate, through methylation/demethylation of the receptors in the case of chemotaxis or sequestration and degradation of sigma factor *σ*^32^ in the case of heat shock response.

Considerable research effort at the interface of Synthetic Biology and Feedback Control Theory focuses on building motifs realising an operation that is indispensable in any feedback mechanism: the comparison of two signals. Examples of such motifs include a switch motif based on integrase and excisionase proteins [Folliard et al., 2017b]; a titration motif [Cuba Samaniego et al., 2016]; an ultrasensitive motif [Samaniego and Franco, 2021,Montefusco et al., 2016]; all these on top of traditional transcriptional feedback, where a transcription factor represses expression of a protein [Becskei and Serrano, 2000]. An important example of ‘subtraction’ is the ‘molecular sequestration’ motif (cf. [Briat et al., 2016]), in which two molecules annihilate each other, thus lowering the functional concentration of both. This encompasses many current implementations of feedback, such as sigma/anti-sigma factors [Annunziata et al., 2017], scaffold/anti-scaffold proteins [Hsiao et al., 2015], mRNA-sRNA interactions [Kelly et al., 2018] and others. In fact, most of the recent research in feedback control for Synthetic Biology is aimed at analysing, realising and applying different versions of this motif [Agrawal et al., 2018,Huang et al., 2018,Aoki et al., 2019,Delalez et al., 2018a,Lillacci et al., 2018]. All of these motifs, however, are usually designed without taking into account the architecture of the process to be controlled - the process is instead used to guide tuning of the controller parameters in order to achieve the desired performance.

Sometimes natural biological systems realise sequestration in a way different to ‘molecular sequestration’. For example, in two-component signal transduction systems (TCSS) having a second response regulator phosphorylated by the same kinase siphons phosphorylation resources away and decreases signal transduction on the main pathway, thus ‘sequestering’ the active kinase signal instead of annihilating specific molecules [Sourjik and Schmitt, 1998]. This, for example, occurs in the *Sinorhizobium meliloti* chemotaxis pathway, where two response regulators CheY_1_ and CheY_2_ are phosphorylated by one kinase CheA. While CheY_2_ is the motor function regulator, CheY_1_ adjusts the sensitivity to the signal [Amin et al., 2014, Sourjik and Schmitt, 1998]. The copy number of the CheY_1_ protein can also be used to modulate the response of the system [Amin et al., 2014]. ‘Signal sequestration’ through competition occurs in other TCSSs; for example it was reported that in the *Escherichia coli* nitrate sensing system, the competition for phosphorylated molecules occurs at three levels: two kinases NarX and NarQ compete for the available ATP molecules, the response regulators NarL and NarP compete for the kinases NarX and NarQ, and the phosphorylated NarL and NarP bind to the same promoters (NarL, however, binds to other promoters as well) [Laub and Goulian, 2007]. In the EnvZ-OmpR system there is also evidence of promoter competition as the phosphorylated OmpR can bind to *pompc* and *pompf* [Laub and Goulian, 2007]. Signal sequestration also appears due to cellular resource depletion such as RNA polymerases and ribosomes needed for transcription and translation initiation, respectively [Ceroni et al., 2015]. In this case, over-expression of one protein can lead to the decrease in expression of another protein, even if they do not directly interact with each other [Qian et al., 2017]. Hence, signal sequestration appears to be one of the natural ways for realising ‘signal comparison’. Synthetic realisations of signal sequestration also exist, for example, in [Potvin-Trottier et al., 2016] another copy of a promoter in an oscillator circuit was introduced in order to fine tune oscillations, and DNA ‘sponges’ were recently considered [Xinyi et al., 2020].

In this paper, we use signal sequestration to develop a new type of feedback regulation, which we term *dichotomous feedback*. This leads to a new framework for designing feedback controllers, leveraging the naturally occurring *signal sequestration motif*. In this feedback, the number of target molecules are decreased by increasing (respectively, decreasing) the rate of a reaction for which the target molecules are the reactants (respectively, products). Since this approach can target existing reactions in the process for implementing the sequestration, this motif can be easier to realise in some situations [Amin et al., 2014]. While different realisations of signal sequestration were studied in the past [Amin et al., 2014,Qian et al., 2017,Potvin-Trottier et al., 2016,Xinyi et al., 2020], in this paper we propose to use signal sequestration to design feedback.

In particular, we propose and analyse theoretically our dichotomous feedback design in a two component signalling system. We realise this feedback architecture by placing a signal sequestration protein downstream of the output protein. The purpose of this protein is to sequester reactions leading to lower concentrations of the phosphorylated response regulator. As a signal sequestration protein, we can use either: (a) a second response regulator, which sequesters the available kinase activity but does not bind to DNA, thus lowering the output protein expression; or (b) a phosphatase for the natural response regulator. It was suggested that in the two response regulator system the second response regulator effectively acts as a phosphatase by sequestering the phosphorylated kinase [Sourjik and Schmitt, 1998]. Hence the second response regulator and the second phosphatase have differences in realisation (affecting different signals), but are similar in effect (reducing the number of phosphorylated natural response regulators). To summarise, we either a) sequester the phosphorylation reaction (by using a second response regulator) or b) enhance the dephosphorylation reaction (using a second phos-photase). Modelling reveals that the designs are, indeed, qualitatively similar and possess comparable properties. The designs differ in terms of feedback strength, which varies depending on the particular TCSSs and signal sequestration proteins.

Lastly, it was also proposed that the competition between two response regulators for a common kinase or between two phosphatases for a common response regulator can be used as means for mitigating cross-talk in natural and synthetic systems [Laub and Goulian, 2007], [Steel et al., 2018]. In this paper we also theoretically verified that our dichotomous feedback architecture reduces the effect of other kinases phosphorylating the natural response regulator. This and other properties are discussed in detail in the rest of the paper.

## Results

### Signal Sequestration Realises Signal Comparison

It is often required that the output of a process, which in the biological context can be the concentration of a particular molecule, follows a desired reference value. In order to achieve this robustly, one must use an engineering feedback control system: the process output needs to be compared with the reference value and based on this ‘error’ quantity, an appropriate input to the process needs to be calculated. Sequestration motifs have been used extensively in Synthetic Biology to realise this operation.

Consider the biochemical process in Figure 1.a, where we want to control the concentration of molecules *Y*: this can be achieved by decreasing the concentration of molecules *U*. In order to do so, the *molecular sequestration motif* introduces molecules *M*, which bind to the molecules *U* and render them inactive, so that the ‘available’ concentration of *U* is decreased. Once sequestration is established, feedback is achieved by producing *M* from *Y*, e.g., using a reaction *Y* → *M*.

**Figure 1:**
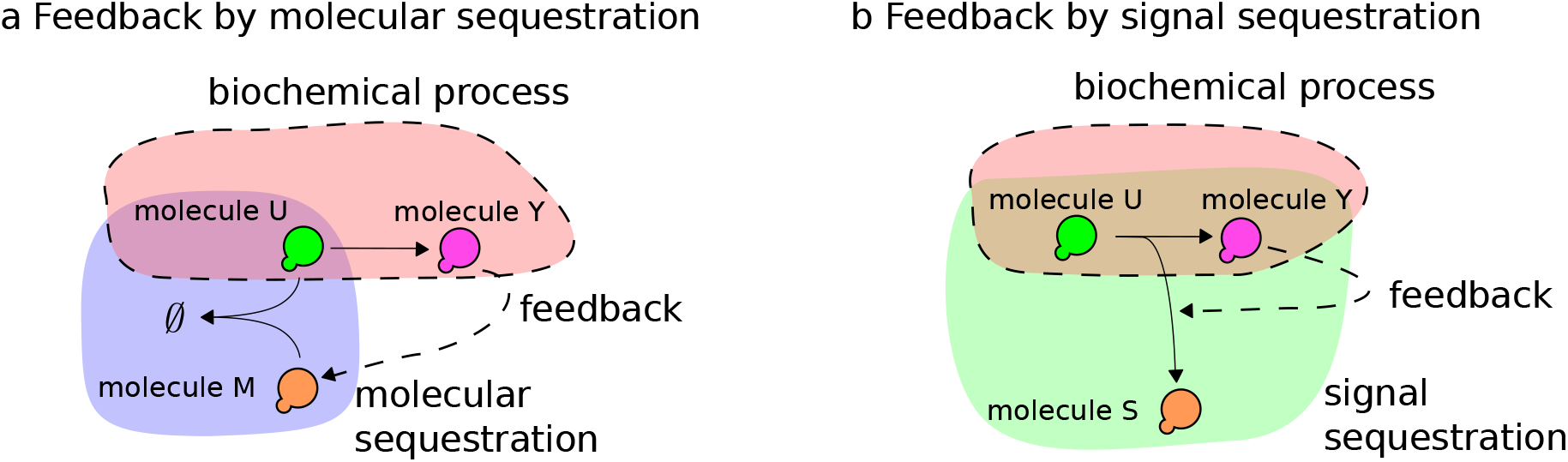
a. Negative feedback by molecular sequestration: Molecule *Y* produces the molecule *M* that binds to molecule *U* thus lowering the concentration of *U*. b. Negative feedback by signal sequestration: Molecule Y increases the reaction rate *U* → *S* thus lowering the concentration of *U*.

Another way of achieving feedback, which we will study at length in this paper, is by sequestering the *signals* involving *U*. In order to decrease the concentration of *U* in the biochemical process depicted in Figure 1.b we add a reaction 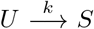 and establish the feedback by affecting the reaction rate *k* through *Y*. In effect, the reaction 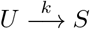 sequesters the signal/flux from *U* to *Y*.

### Signal Sequestration in Two-Component Systems

The concept of signal sequestration is easier to explain using a motivating example: Signal transduction in two-component signalling systems (TCSS). In these systems signalling occurs by transferring phosphoryl groups. First, a histidine kinase (HK) is autophosphorylated, e.g., in the presence of a chemical inducer (I), and thereafter a response regulator (RR) receives the phosphoryl group from the kinase. The phosphorylated response regulator (RR_p_) then acts as a transcription factor activating (or in some cases inhibiting) an output protein’s (Output) expression. A chemical reaction model describing such a two-component system is as follows:

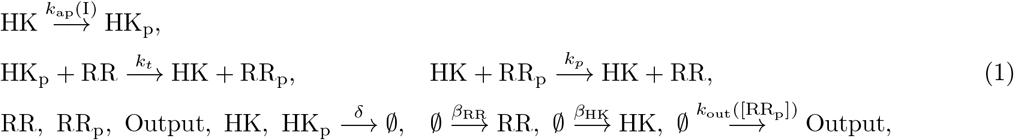

where the subscript ·_p_ denotes the phosphorylated form of a protein, *k*_ap_(I), *k_p_*, *k_t_* are histidine kinase autophosphorylation, response regulator phosphorylation and response regulator dephosphorylation rates, respectively; *δ* is the dilution rate; and *β*_HK_, *β*_RR_ are the HK and RR production rates, respectively. We assumed that the autophosphorylation rate *k*_ap_(I) depends on the inducer concentration as a Michaelis-Menten function, while the output expression initiation *k*_out_([RR_p_]) depends on the response regulator as a Hill function, as previously suggested [Chang et al., 2013]:

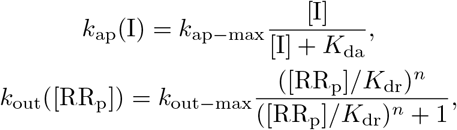

where *k*_ap–max_ is the maximal autophosphorylation rate, *K*_da_ is the inducer dissociation constant, *k*_out–max_ is the maximum production rate of the output protein, *K*_dr_ is the dissociation constant, and *n* is the cooperativity coefficient.

Signalling systems like this tend to exhibit a particularly simple and robust behaviour as discussed in detail in Section B.3 in SI. For example, there is typically a unique asymptotically stable steady state for a fixed inducer concentration, which increases monotonically with increased inducer concentrations. Furthermore, the response curve to the inducer is a Hill function, hence the response saturates for a large inducer value. For many systems the Hill functions are very steep, which effectively gives two modes: on (for large inducer concentrations) and off (for small inducer concentrations). This necessarily implies that achieving intermediate response values is difficult in the natural system, but is possible through feedback reducing the concentration of RR_p_. This, in effect, exploits the limited number of phosphoryl groups in the system. There are two different signal sequestration motifs that can achieve this: Sequestration of phosphorylation of RR_p_; and enhancement of dephosphorylation of RR_p_, as depicted in Figure 2.

**Figure 2:**
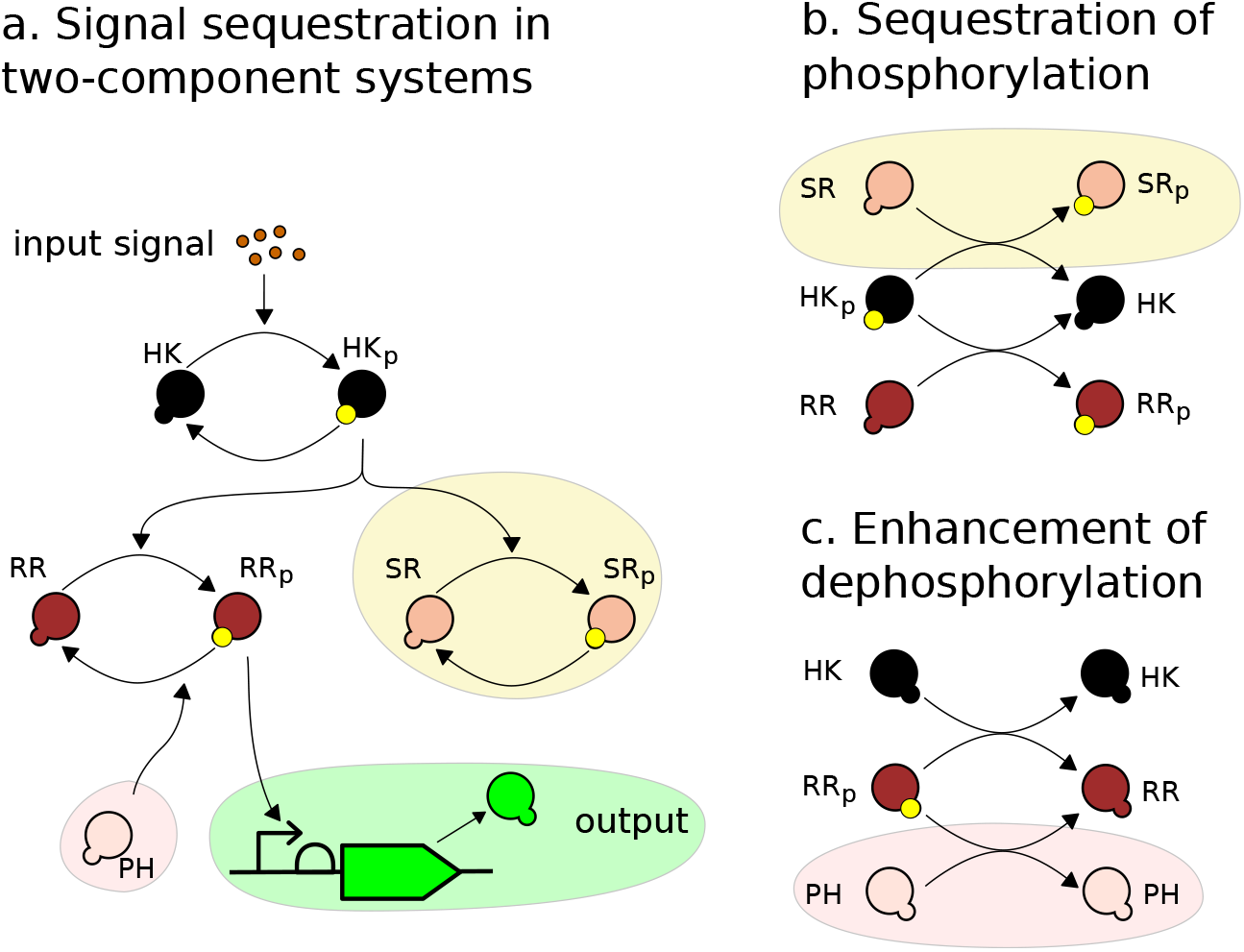
a. A schematic depiction of two signal sequestration motifs in a two-component system. We do not depict the enzymatic nature of the dephosphorylation reaction of the histidine kinase, the response regulator and the sequestration proteins for visualisation purposes. b. Phosphorylation sequestration. As the signal is transferred through the histidine kinase it is sequestered by the phosphorylation of the sequestration protein thus fewer response regulators are being phosphorylated decreasing the output signal. c. Dephosphorylation enhancement. A phosphatase is dephosphorylating the response regulator in addition to the action of histidine kinase thus fewer response regulators are phosphorylated.

In *phosphorylation sequestration*, a second response regulator, the protein SR, is also phospho-rylated and dephosphorylated by the same histidine kinase as the response regulator, but does not transcriptionally activate the output protein, thus sequestering the phosphotransfer flux. We modelled the phosphorylation sequestration using the following chemical reactions:

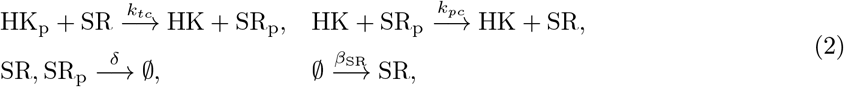

where *β*_SR_ is the production rate of SR.

The second way of achieving feedback is by *enhancing dephosphorylation*, realised by producing a phosphatase (PH) to dephosphorylate the response regulator – this is modelled using the reactions:

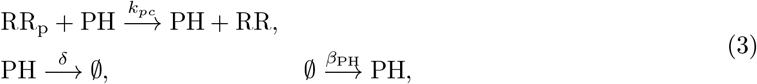

where *β*_PH_ is PH production rate.

Assuming these chemical reactions and mass-action kinetics, we can obtain the following differential equation model:

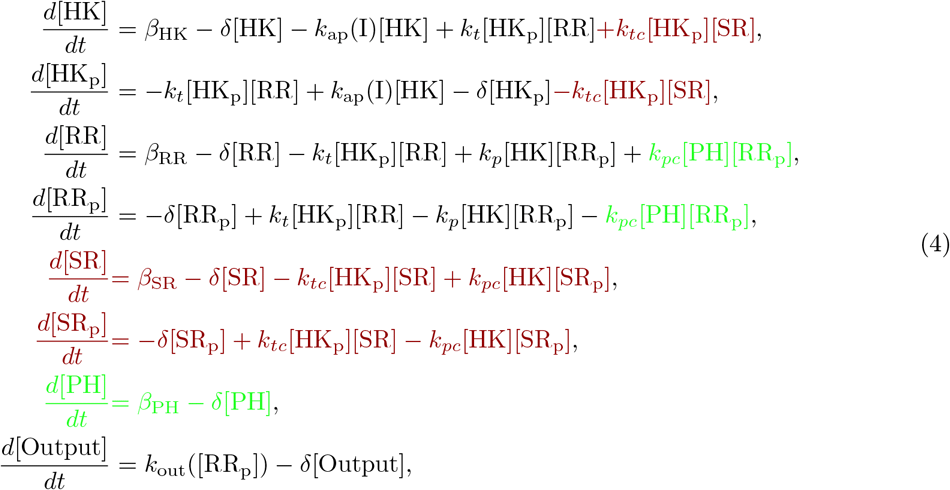

where we mark phosphorylation sequestration and dephosphorylation enhancement reactions in burgundy and green, respectively, and depict this model schematically in Figure 2.a. These sequestration mechanisms are ‘dual’ to each other - in the sense they operate on the two sides of the phosphorylation process while having similar expressions for their steady-state response, after a suitable choice of parameters - and therefore their behaviour is qualitatively similar (see Figures 2.b and 2.c for a depiction of this ‘duality’). In order to show this we derived the steady-state models for these motifs, summarised in the following Proposition; the proof can be found in Sections B.4 and C.3 of the SI).

#### Proposition 1

*Let* RR_tot_ = *β*_RR_/*δ*, SR_tot_ = *β*_SR_/*δ*, HK_tot_ = *β*_HK_/*δ*, *and* PH_tot_ = *β*_PH_/*δ*.

1. *Consider the phosphorylation sequestration motif* (*i.e*., *the system* (4) *with β*_PH_ = 0) *under the following assumptions*

a. *The total concentration of the response regulators is much larger than the concentration of the phosphorylated proteins:* [RR_p_] ≪ [RR_sum_], [SR_p_] ≪ [SR_sum_]
b. *The following holds*:

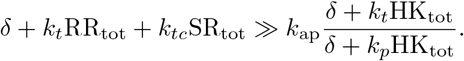 *The the steady state of* [RR_p_] *can be approximated as follows*:

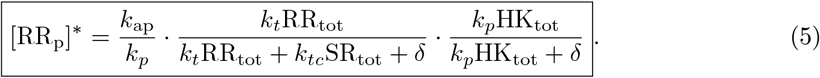
2. *Consider the dephosphorylation enhacement motif* (*i.e*., *the system* (4) *with β*_SR_ = 0). *If the following holds*:

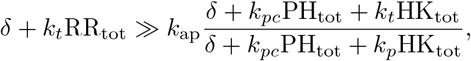

*then the steady state of* [RR_p_] *can be approximated as follows*:

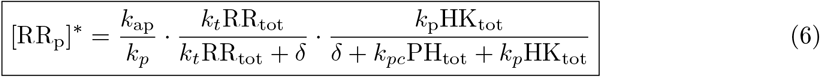

While the realisation of these sequestration motifs is quite different, in terms of the steady-states these differ only in the magnitude of RR_p_ output, which showcases their ‘duality’.

While dephosphorylation enhancement is one of the obvious solutions to decrease the concentration of RR_p_ [Chang et al., 2013, Jones et al., 2021], phosphorylation sequestration is not. Furthermore, the phosphorylation sequestration may be easier to realise in some systems - indeed, engineering a second response regulator is experimentally feasible [Amin et al., 2014] and is particularly valuable if there are no known phosphatases for a particular response regulator.

### Realising feedback through signal sequestration: Dichotomous Feedback

Realising negative feedback using these sequestration motifs can be achieved by placing the *sr* or the *ph* genes under the same promoter as the output gene meaning that their production rates (*β*_SR_, *β*_PH_) depend on [RR_p_]. Both proteins SR and PH sequester phosphoryl groups: While the phosphorylated protein SR keeps the phosphoryl groups attached, the phosphatase PH releases the group into the pool of free phosphoryl groups, but HK would need to be phosphorylated again to access these groups. We will refer to the feedback mechanisms realised with phosphorylation sequestration and dephosphorylation enhancement as *dichotomous feedback* for signalling processes (see Figure 3). In what follows we discuss phosphorylation sequestration in detail since the dephosphorylation enhancement motif exhibits many similar properties; a detailed comparison of the feedback systems can be found in Section D.3 in the SI.

**Figure 3:**
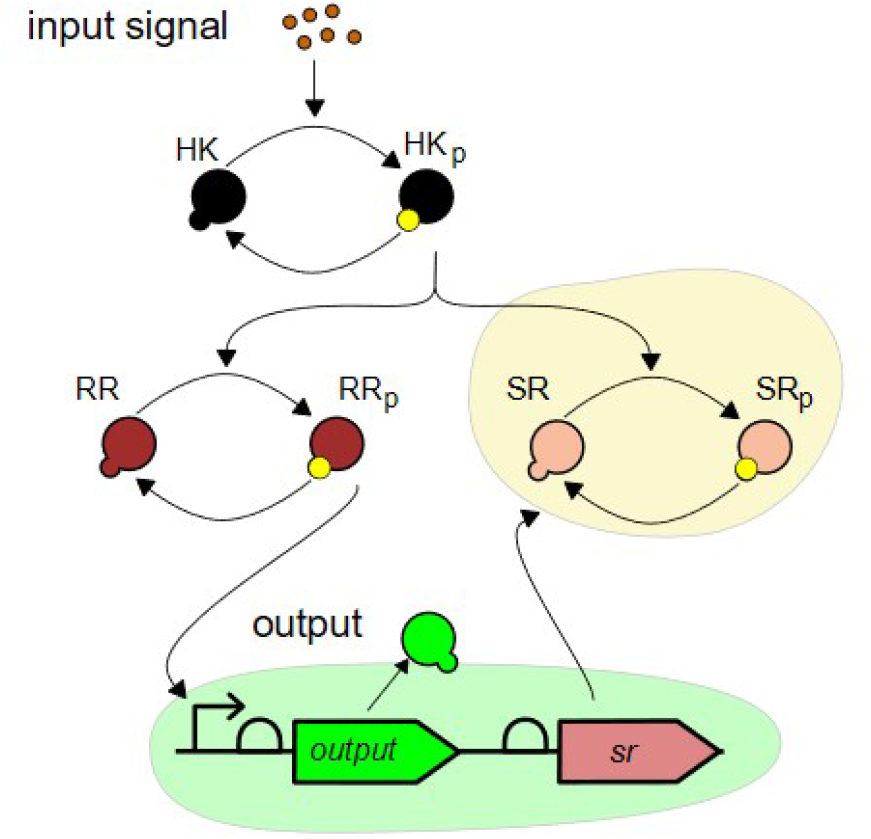
Design of the dichotomous feedback system using phosphorylation sequestration via second response regulator.

We can vary the feedback strength *P* by, e.g., placing an *X*% efficient terminator between the output and the sequestering proteins [Chen et al., 2013] or by varying the strength of ribosome binding site for the sequestering protein [iGEM, 2021]. We note that other possibilities of tuning the feedback strength are possible [Arpino et al., 2013, Ang et al., 2013], for example, one can use small RNAs similarly to [Delalez et al., 2018b]). We therefore adjust our model by considering the following reaction of the sequestration protein expression:

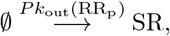

where *P* is the feedback strength. In the differential equation (4), we would replace the constant *β*_SR_ with a variable production rate:

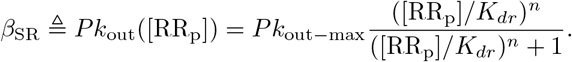

We discuss mathematical properties of this model in what follows, while the model parameters were taken from Table D.1 in SI. The dichotomous feedback based on de-phosphorylation enhancement is designed similarly, see Section C in SI.

### Mathematical Properties of the Dichotomous Feedback

Verifying stability of the signalling cascades is well studied in the literature and our sequestration of phosphorylation motif can be treated by existing methods (see, e.g. [Steel et al., 2018, Steel et al.,2019]). First we will slightly simplify the system

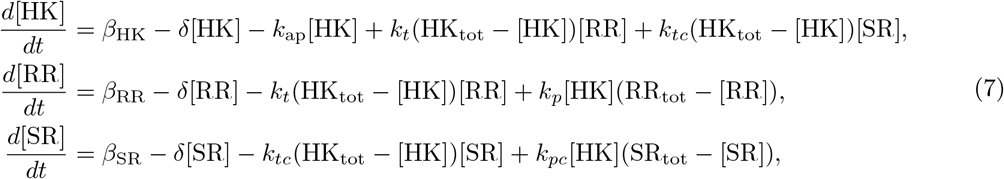

where HK_tot_ = *β*_HK_/*δ*, RR = *β*_RR_/*δ* and SR_tot_ = *β*_SR_/*δ*. We will the study the system properties on the set 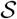:

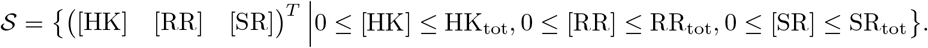

The key to our further discussion is the ability of adding sequestration reactions *without* breaking the major structural mathematical property of the signal transduction pathway — monotonicity. We present a short mathematical introduction to monotone systems in Section A.2 in the Supplementary Information, but also refer the reader to [Angeli and Sontag, 2003, Hirsch and Smith, 2006] for more details. While the theory of monotone systems can be involved, the intuition behind monotone systems is quite straightforward. In general, it is hard to compare the trajectories *φ*(*t*, *x*^0^) of a system knowing only the initial conditions *x*^0^. However, for a monotone system a simple relation holds: Let *z_i_* denote the *i*-th component of the vector *z*, if the system is monotone then if 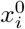 is larger than 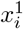 for all *i* then 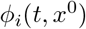 is larger than 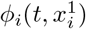 for all *t* > 0. This simple property leads to a number of powerful results in the literature and in our case to the following result:

#### Proposition 2

*Consider the phosphorylation sequestration motif (i.e., system* (7)) *with nonnegative parameters. Then*

1. *Trajectories of the system* (7) *cannot leave the set* 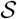;
2. *Consider the natural two component system, i.e., the system* (7) *with β*_SR_ = 0. *The equilibrium* [HK]^*^, [RR]^*^ *of this system is locally asymptotically stable if*

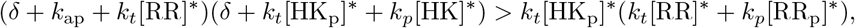

*where* [HK_p_]^*^ = HK_tot_ — [HK]^*^, [RR_p_]^*^ = RR_tot_ — [RR]^*^. *In particular, this condition is satisfied if*:

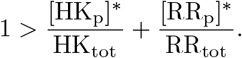
3. *The system* (7) *is not chaotic and cannot have stable limit cycles*;
4. *Consider two trajectories of the system* (7) *φ*_1_(*t*, *x*_1_) *with φ*_2_(*t*, *x*_2_) *with x*_1_ = [HK] [RR] [SR] = (0 0 0) *and x*_2_ = ([HK] [RR] [SR]) = (HK_tot_ RR_tot_ SR_tot_). *If both these trajectories converge to the same point x*^*^ *as t grows to infinity, then x*^*^ *is globally attractive in* 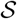;
5. *If the system* (7) *has a globally asymptotically stable in* 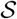 *equilibrium for* 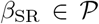, *where* 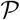 *is an interval in* 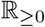, *then the steady-state concentration of* [RR] *increases monotonically with* 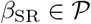.

In order to determine closed-loop stability for a specific set of parameters, we can use a small-gain theorem for monotone systems from [Angeli et al., 2014]. We reproduce the result as Theorem S2 in Section A.2 in Supplementary Information. In particular, to verify stability using this small gain result, we consider the following system:

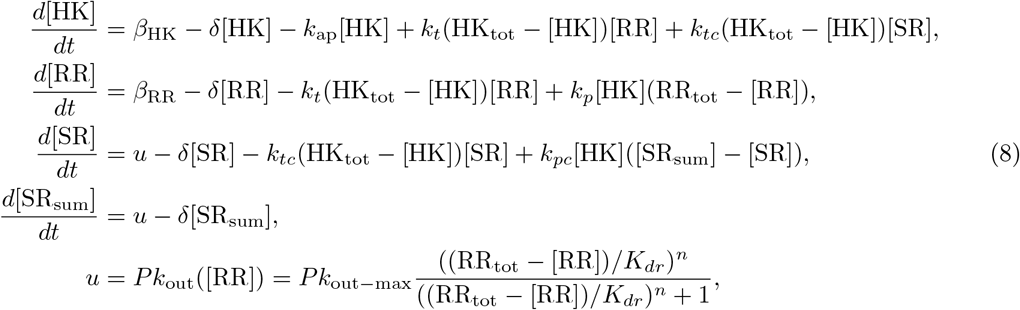

where we treat *β*_SR_ as an input *u* to the signal sequestration motif (the open-loop system (4)) with an output *y* = RR_tot_ — [RR]. Closing the loop is performed by the relation *β*_SR_ = *Pk*_out_ ([RR]).

Analysis of the dephosphorylation enhancement motif is similar.

### Dichotomous Feedback Shapes the Signal Response

We consider models of the wild-type two-component system, the dichotomous feedback architecture with various feedback strengths *P*, and an open-loop sequestration architecture in which the protein SR is under an inducible promoter and hence we can vary the total steady-state concentrations of the protein SR computed as SR_tot_ = *β*_SR_/*δ*. The design of the system is schematically depicted in Figures 4.a and 4.b. The dose-response curves depicted in Figure 4c show that the dichotomous feedback architecture allows us to achieve a range of maximum response values by varying the feed-back strength. The open-loop sequestration architecture offers a graded response to the inducer for a particular value of SR_tot_, while the dichotomous feedback offers a sigmoidal response. Therefore, the open-loop response is less robust to variations in the inducer concentrations in comparison to the dichotomous feedback architecture at large induction levels. Note that the maximum value in the sigmoidal response of the dichotomous feedback depends on the terminator efficiency and is hence tunable allowing to achieve intermediate response values in a robust manner. This is not possible in the wild-type signalling system. To summarise, only the dichotomous feedback can achieve intermediate response values at large induction levels in a robust fashion using different feedback strengths. We performed simulations for different phosphorylation and dephosphorylation rates in Section D.3, which yielded similar results. The results for the phosphatase feedback architecture (Figure 2.c), also presented in Section D.3, indicate that the feedback mechanisms share these properties.

**Figure 4:**
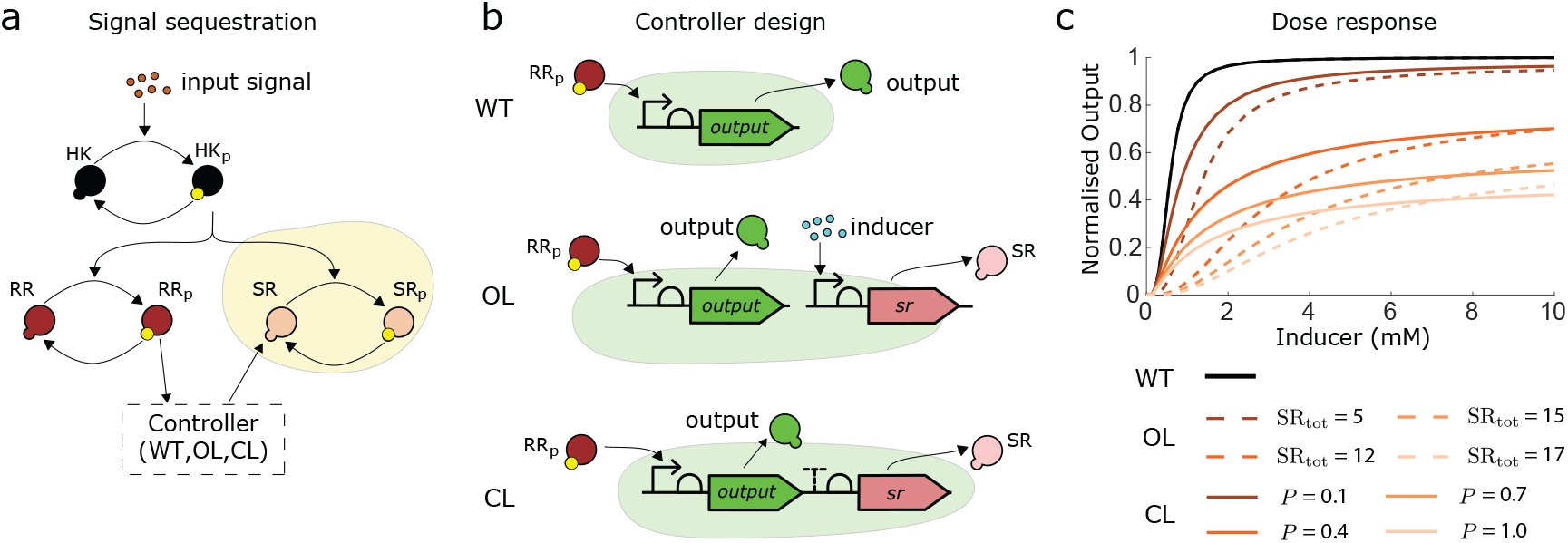
a. Interactions in the phosphorylation sequestration system. b. Design of the wild-type (WT), the open-loop phosphorylation sequestration motif (OL) and the closed loop dichotomous feedback (CL) systems. c. Dose-response curves of the wild-type, the phosphorylation sequestration motif (open-loop) and the dichotomous feedback (closed-loop) systems. Closing the loop allows achieving intermediate values of the output protein expression by tuning the feedback strength.

### Dichotomous Feedback Attenuates Intrinsic Noise

We estimated the intrinsic noise properties of the dichotomous feedback and the wild-type systems using the Linear Noise Approximation (LNA) modelling framework, which models a chemical reaction as an evolution of the Gaussian distribution with the mean modelled by a nonlinear ordinary differential equation and the variance modelled by a linear matrix differential equation. We describe this framework in detail Section A.4.2 in SI. As a metric for estimating noise levels we took the coefficient of variation, which is defined as follows:

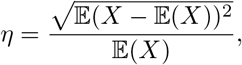

where 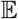 stands for the mean of the random variable *X*. In order to verify the accuracy of our computations, we also computed the coefficient of variation at time *t* = 1000 [min] for specific inducer concentrations and feedback strengths using 10^5^ Gillespie stochastic simulations. We plot the estimated coefficient of variation of the wild-type and the dichotomous feedback systems against the mean output response in Figure 5.a. The dichotomous feedback architecture allows us to reach various mean values of the output protein while reducing intrinsic noise. Interestingly, increasing the feedback strength decreases the coefficient of variation, however, it is known that in many feedback systems increasing the feedback strength can potentially increase intrinsic noise [Singh and Hespanha, 2009,Oyarzun et al.,2015]. Similar results apply to a range of phosphorylation and de-phosphorylation rates, as well as the dephosphorylation enhancement motif (see Section D.3 in SI). As LNA provided quantitatively similar coefficient of variation computations to stochastic simulations we use LNA to perform this analysis.

**Figure 5:**
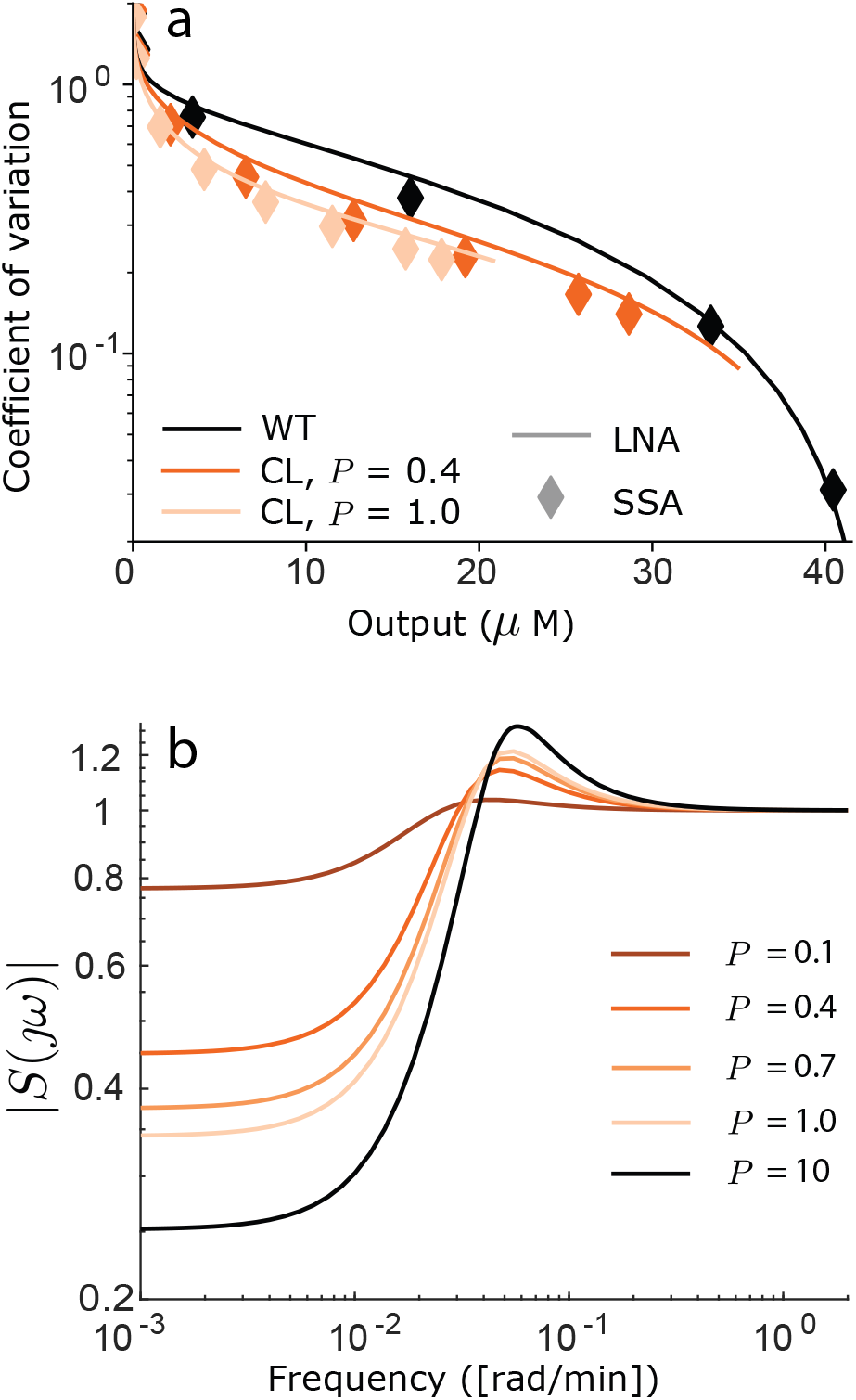
a. The coefficient of variation of the wild-type and the closed-loop systems against the mean output protein concentrations. We computed the coefficient of variation using the Linear Noise Approximation (LNA, see Section A.4.2 in SI) and using 10^5^ runs of the Gillespie stochastic simulation algorithm (SSA) at time *t* = 1000 [min]. The plot suggests that the intrinsic noise levels are reduced in the closed-loop in comparison with the wild-type system. b. The magnitude of the sensitivity function of the closed-loop systems for [*I*] = 2 [mM]. The shape and the values of the magnitude of the frequency response suggest that the closed-loop system is robust towards perturbations.

### Dichotomous Feedback Implements a Robust Controller

We then studied the robustness of our feedback system using feedback control theoretic methods, which we describe in Section A.3 in SI. As we study the behaviour of the system around a stable steady state we resort to a local analysis. The local analysis is performed by linearising the system around the steady state, which is equivalent to studying a nonlinear system locally under some mild assumptions [Olsman et al., 2019]. We then use frequency domain analysis and transfer functions, which are equivalent operator representations of a linear system. In particular, we will consider a specific transfer function, the sensitivity function *S*(*jω*), where *ω* is a real-valued frequency describing the frequency of the input. The objective is to calculate the effect that a disturbance/perturbation will have on the properties of a closed loop system. To do this, the closed-loop system is broken down to a process (in our case a two-component system) and a controller (in our case the sequestration motif). The process model is as follows:

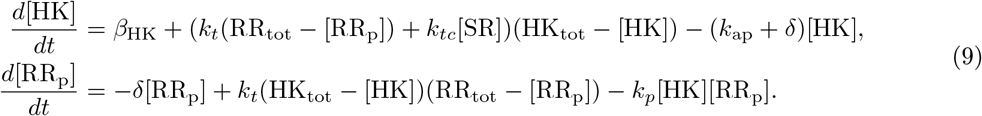

while the model for the controller is:

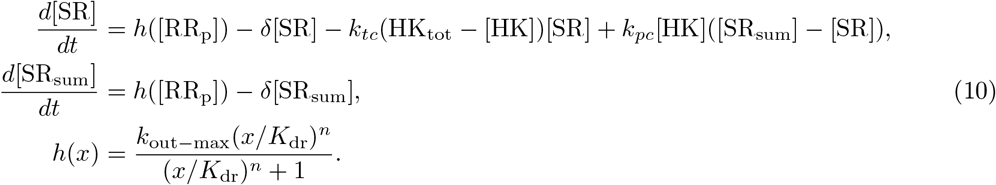

For the process model we assume that [HK_sum_] = HK_tot_ and [RR_sum_] = RR_tot_, since we perform the analysis around the steady-state. Hence the process’ states are [HK], [RR_p_], while controllers are [SR], [SR_sum_]. The process has one external input – the inducer concentration entering the equation through the autophosphorylation rate *k*_ap_, which we denote as *z*. The process also has a controlled input [SR] (denoted as *u*) and the outputs [HK] (*y*_1_) and [RR_p_] (*y*_2_), while the controller has the output [SR] and the inputs [HK] and [RR_p_]. Now we can compute the transfer functions of the linearised model and the linearised controller as described in Section A.3 in SI. The inputs and outputs are related using the following equation

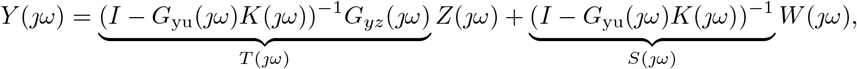

where *G*_yu_, *G*_yz_ are transfer functions from *U*(*jω*) to *Y*(*jω*) and from *Z*(*jω*) to *Y*(*jω*), respectively; *W*(*jω*) models the process disturbances (such as, modelling errors), *Z*(*jω*) models the external inputs and *Y*(*jω*) models the output of the process. The setting is depicted in Figure 6, which we use in this paper.

**Figure 6:**
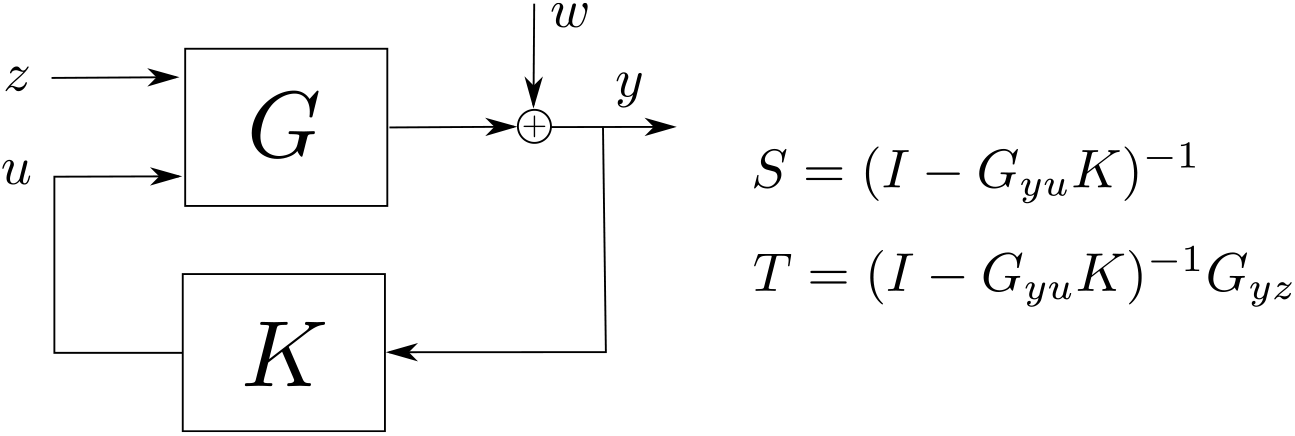
Block diagram of the process *G* to be controlled and the controller *K* feedback loop used in this paper. The signal *z* signifies the external inputs, *y* the measured output, *w* are the process disturbances (e.g., modelling errors or external perturbations) and *u* is the control signal computed by the controller *K*. The significance of the functions *S* and *T* is discussed in the text below.

The function *T*(*jω*) describes the behaviour of the system with respect to the external inputs *Z*(*jω*), i.e., the concentration of the inducer. The function *S*(*jω*) describes the contribution of the disturbance *W*(*jω*) to the output, which ideally should either attenuate the disturbances (i.e., the magnitude of *S*(*jω*) for 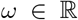 is close to zero) or at least not amplify them (i.e., the magnitude of *S*(*jω*) for 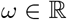 is close to one). In control engineering, the rule-of-thumb is to design the controller *K* so that 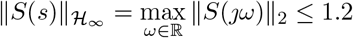 [Astrom and Murray, 2010]. Low magnitude values of the sensitivity function for small frequencies are also preferable. For example, if the magnitude at zero frequency is equal to zero, then the feedback architecture achieves perfect adaptation [Astrom and Murray, 2010].

We plot the magnitude of the sensitivity function for various feedback strengths in Figure 5.b. The maximal magnitude of the sensitivity function for the inducer concentrations [0.5, 1, 2, 5, 10] and feedback strengths [0.1, 0.4, 0.7, 1] was equal to 1.2124 achieving the maximum for [*I*] = 2 [mM] and *P* = 1. These computational results imply that the dichotomous feedback architecture is robust to disturbances and theoretically can allow for close-to-perfect adaptation for appropriately large feedback strengths. We performed simulations for different phosphorylation and dephosphorylation rates in Section D.3 in SI, which yielded similar results. In the case of the phosphotase we also assume that [HK_sum_] = HK_tot_ and [RR_sum_] = RR_tot_ and separate the model into the process and the controller. The process equations are:

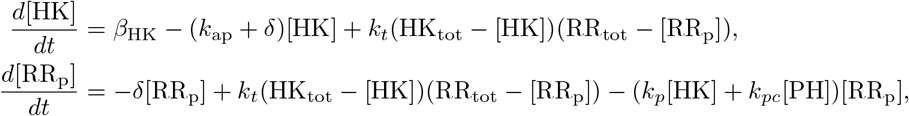

where we treat [PH] as the input and the phosphorylated RR as the output. The controller equations are

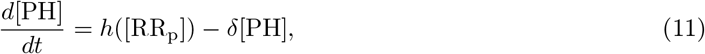

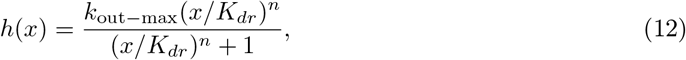

where we treat RR_p_ as an input. It is clear that the phosphotase feedback is simpler to design and analyse since the controller has just one state. The phosphotase feedback appears to have a much larger feedback strength due to its direct action on kinase fluxes. However, as our simulations (presented in Section D.3 in SI) indicate the behaviour of both feedbacks is qualitatevely similar otherwise. Furthermore, these results indicate that the feedback mechanisms share the robustness property.

### Differences Between Molecular and Signal Sequestration Motifs

While two-component signalling cascades are important examples of signal sequestration, we now discuss whether signal sequestration is a different design mechanism from molecular sequestration, and whether there are other examples of signal sequestration motifs in natural and synthetic systems.

In their simplest implementations, molecular and signal sequestration can be achieved in a similar way, especially if the sequestration is achieved through the aid of another molecule. Their differences become evident in how the two can be implemented in practice. In fact, signal sequestration is a straightforward choice for the designer if in the reaction *U* → *Y* the reactant *U* is multi-functional, for example, RNA polymerase (RNAP) or a transcription factor enhancing transcription, a ribosome or an activating sRNA [Chappell et al., 2015] initiating translation as depicted in Figure 7. More complex designs involving riboswitches [Folliard et al., 2017a], [Breaker, 2012], [Caron et al., 2012] can also be exploited to create signal sequestration. Translational and transcriptional signal sequestration motifs have been considered indirectly in the context of resource sharing (specifically, RNAP and ribosomes) in synthetic circuits [Qian et al., 2017]. A transcription sequestration motif based on the transcription factor was suggested in [Potvin-Trottier et al., 2016] in order to tune the oscillations in the synthetic repressilator. To close the loop, one needs to use the output *Y* to produce the DNA fragment containing the gene for *S* or the mRNA containing the mRNA of *S* and translationally activated by *U*. Producing the DNA fragments is possible using a reverse transcriptase [Farzadfard and Lu, 2014], for example, while producing new mRNA strands can be performed using standard transcriptionally activated promoters.

**Figure 7:**
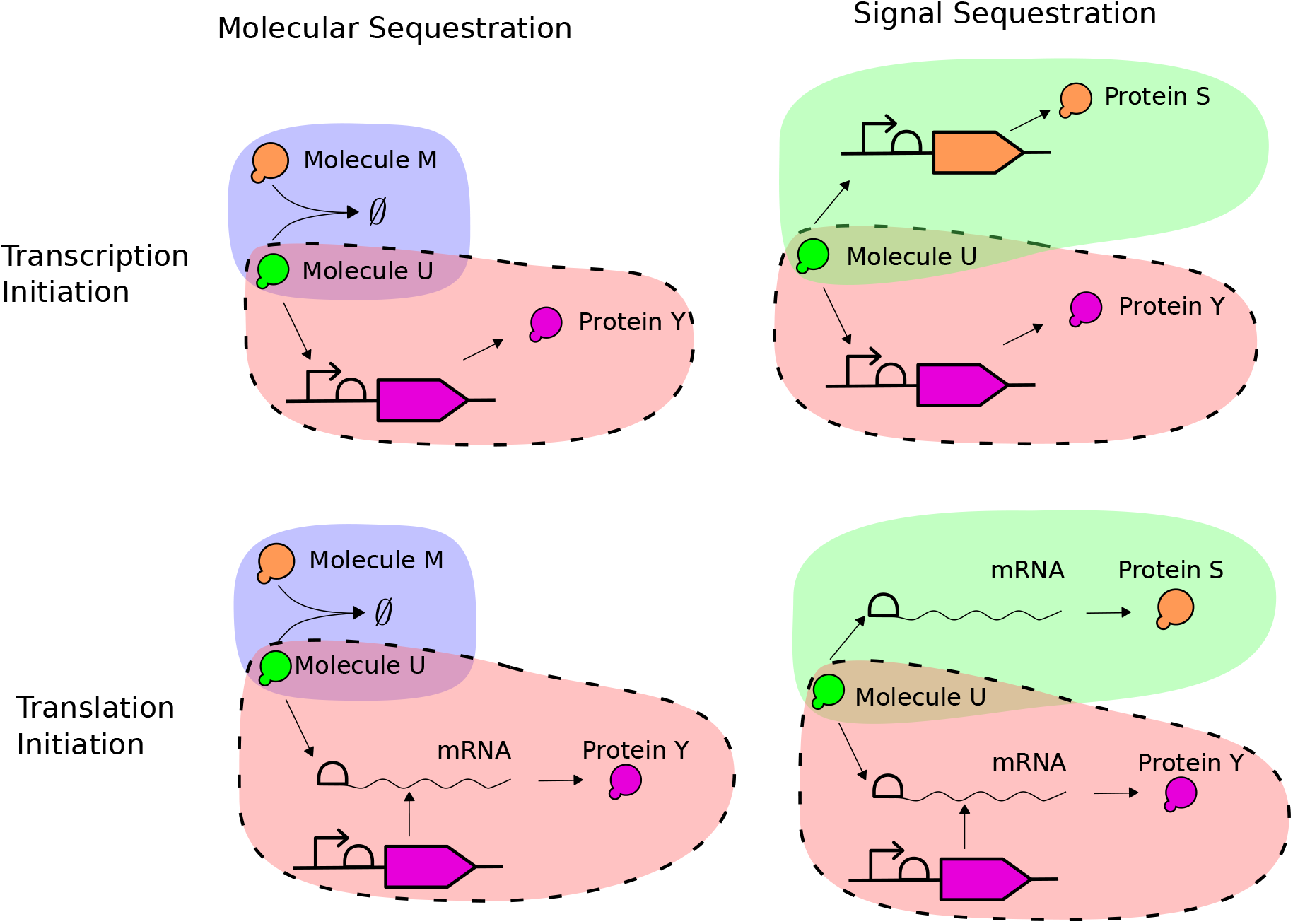
Molecular and signal sequestration for transcription and translation initiation reactions. Shaded areas signify molecular sequestration (blue), signal sequestration (green) motifs and the process to be controlled (red). Possible realisations of negative feedback are not depicted for visualisation purposes. In molecular sequestration, molecule *M* annihilates the molecule *U*, while in the signal sequestration the reaction involving the molecule *U* is reproduced thus siphoning the signal to produce the protein *S*. In molecular sequestration motifs, negative feedback can be realised by producing the molecules *M* from the molecules *Y*, e.g., using a reaction *Y* → *M*. In signal sequestration motifs, it is necessary to produce the DNA fragment with the gene *S* and the promoter in transcription initiation, and the mRNA strand containing a ribosome binding site and the mRNA of *S* in translation initiation.

We use the modelling results from [Qian et al., 2017] in order to study the transcriptional and translational signal sequestration motifs. In particular, a simplified differential equation model describing the concentration of proteins *Y* and *S* can be written as follows:

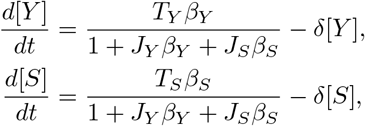

where *δ* is the dilution/degradation rate of the proteins *Y* and *S*, and *β_Y_*, *β_S_* are the expression initiation rates, *T_Y_*, *T_S_* are the baseline expression rates and *J_Y_*, *J_S_* are measures of resource usage (here the resource in question is the shared pool of *U*). Resource sequestration can be better understood using the steady-state model for *Y*:

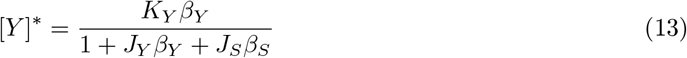

Here *K_Y_* = *T_Y_*/*δ* is a lumped parameter signifying basal expression. Note the similarities in the shape of the steady-state formula (13) and the formulae in (5) and (6). This again highlights similarities between the architectures. The production of *S* sequesters the transcription or translation initiation signal thus lowering the steady-state values of *Y. Dichotomous feedback* can then be realised by linking the production rate *β_S_* with the concentration [*Y*]. If there is an abundance of resources then *J_Y_* and *J_S_* are small and close to zero and *Y* becomes approximately equal to *K_Y_β_Y_*. In this case, the feedback strength through the signal sequestration is very weak and the protein *Y* is expressed in almost normal operation. If there are no free molecules *U* (*J_Y_* and *J_S_* are large), then the signal sequestration feedback becomes much stronger as the molecules *U* must be shared between the processes. Therefore, the signal sequestration feedback can potentially be used in resource-limited systems.

### Dichotomous Feedback is a Different Feedback Mechanism

As the sequestration response regulator binds to the kinase, we compare our dichotomous feedback to two feedback mechanisms depicted in Figures 8.a and 8.b:

a. Molecular sequestration based feedback annihilating the histidine kinase (phosphorylated or not).
b. Transcriptional feedback repressing the production of the histidine kinase.

**Figure 8:**
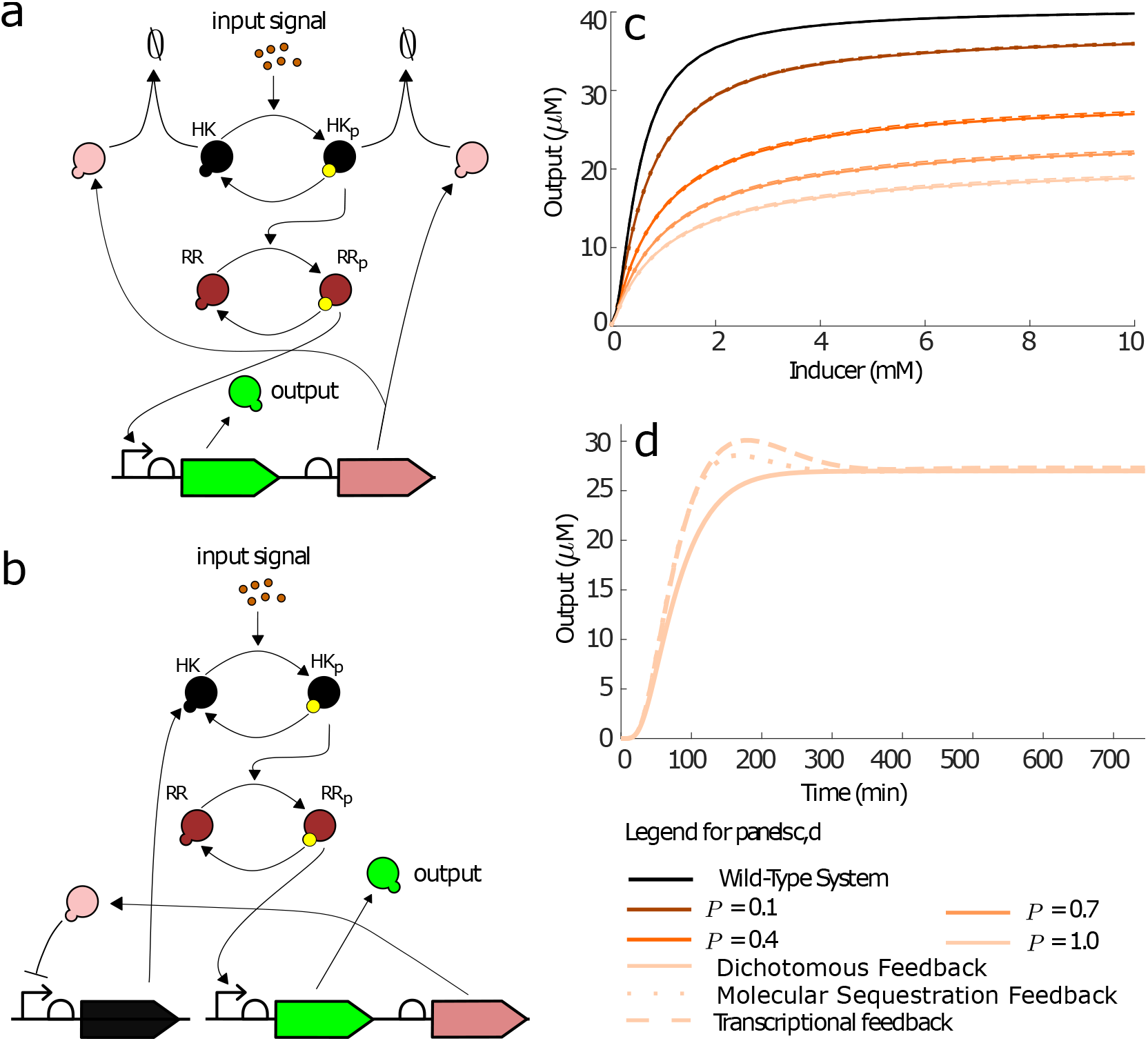
a. Design of the molecular sequestration-based feedback controlling the output. The feedback is realised by a protein annihilating the histidine kinase in its unphosphorylated and phosphorylated forms. b. Design of the transcriptional feedback controlling the output. The feedback is realised by a protein repressing the expression of the histidine kinase HK. c. Dose response curves for different implementation of feedbacks (curves for the same value of *P* overlap). The qualitative behaviour of the dose-response curves is similar for all feedbacks. We picked the strength of the feedback loops so that dose-response curves approximately match. d. Time response of the feedbacks with *P* =1 and [*I*] = 10 [mM]. The transcriptional feedback is the slowest feedback, since the transcription factor first has to be expressed, then lowers the production rate of the histidine kinase, the decrease in the concentration occurs through dilution. The responses for the transcriptional and molecular sequestration based feedbacks have an undesirable overshoot.

We selected parameter values for these architectures so that their steady-state responses match with the dichotomous feedback architecture (see Figure 8.c). The modelling details and parameter values are presented in Section D.4.

The time evolution of the output of each the feedback architecture to a step input (depicted in Figure 8.d) shows that the time responses for the transcriptional and molecular sequestration based feedback architectures have an overshoot in the output protein expression, which is not a desirable property for feedback control. The dichotomous feedback architecture, on the other hand, does not exhibit an overshoot. The overshoot occurs since the transcriptional and molecular sequestration based feedback architectures reduce the total concentration of the histidine kinase forcing the two-component system to adjust to the new setting. As the dichotomous feedback merely acquires the phosphoryl group from the histidine kinase no such adjustment is required. The simulations depicted in Figure S6.a and Figure S6.b in SI also suggest that both transcriptional and molecular sequestration based feedback architectures are less robust to disturbances as the sensitivity function magnitudes are larger than in the dichotomous feedback. Furthermore, computing the magnitude of the sensitivity functions for inducer concentrations from [0.1, 1, 2, 5, 10](mM) and *P* = [0.1, 0.4, 0.7, 1] yielded maximum values of 1.6242 and 1.5124 for the transcriptional and molecular sequestration based feedbacks, respectively, implying a significant sensitivity to some disturbances. If the molecular sequestration based feedback is realised by sequestering *only* the phosphorylated HK or the phosphorylated RR then its sensitivity functions are almost identical to those of the dichotomous feedback. This indicates that the molecular sequestration based feedback architecture can be more sensitive to system structure than the dichotomous feedback, which can take structural properties of a system into account by enhancing or sequestering existing system fluxes.

### Dichotomous Feedback Reduces Crosstalk

Signalling cascades are often interconnected with other signalling cascades. For instance, a response regulator can be phosphorylated by a number of different kinases (see [Laub and Goulian, 2007] for specific examples). We considered the situation in which another histidine kinase *X* is phosphorylated through an unknown mechanism and passes the phosphoryl group to the response regulator RR thus increasing the response even in the absence of the inducer I, as a way to emulate crosstalk. If our sequestration protein is also phosphorylated by the kinase *X*, then the dichotomous feedback architecture can reduce the crosstalk with the histidine kinase *X*. In Figure 9 we depict the architecture of our crosstalk system and the dose-response curves of the wild-type and the closed-loop systems. We chose the constant phosphorylation rate for *X* equal to 0.08 [1/min] and assumed that *X* phosphorylates and dephosphorylates RR and SR at the same rates as HK. Numerical simulations suggest that the signalling feedback architecture lowers the basal response of the system (Figure 9.c), which arises because of the crosstalk with *X*. At the same time, the system still responds to the increase in the inducer concentration. Only when a basal level (independent of [RR_p_] and [I]) of expression was added to the output promoter did basal expression stay the same for all feedback strengths, see Figure S7.b in SI. The molecular sequestration based and transcriptional feedback architectures have minimal effect on the basal expression and hence do not have any effect on crosstalk, see Figure S7.a in SI. The simulation results for the phosphatase feedback architecture were indistinguishable from the results in Figure 9 for the chosen parameter values.

**Figure 9:**
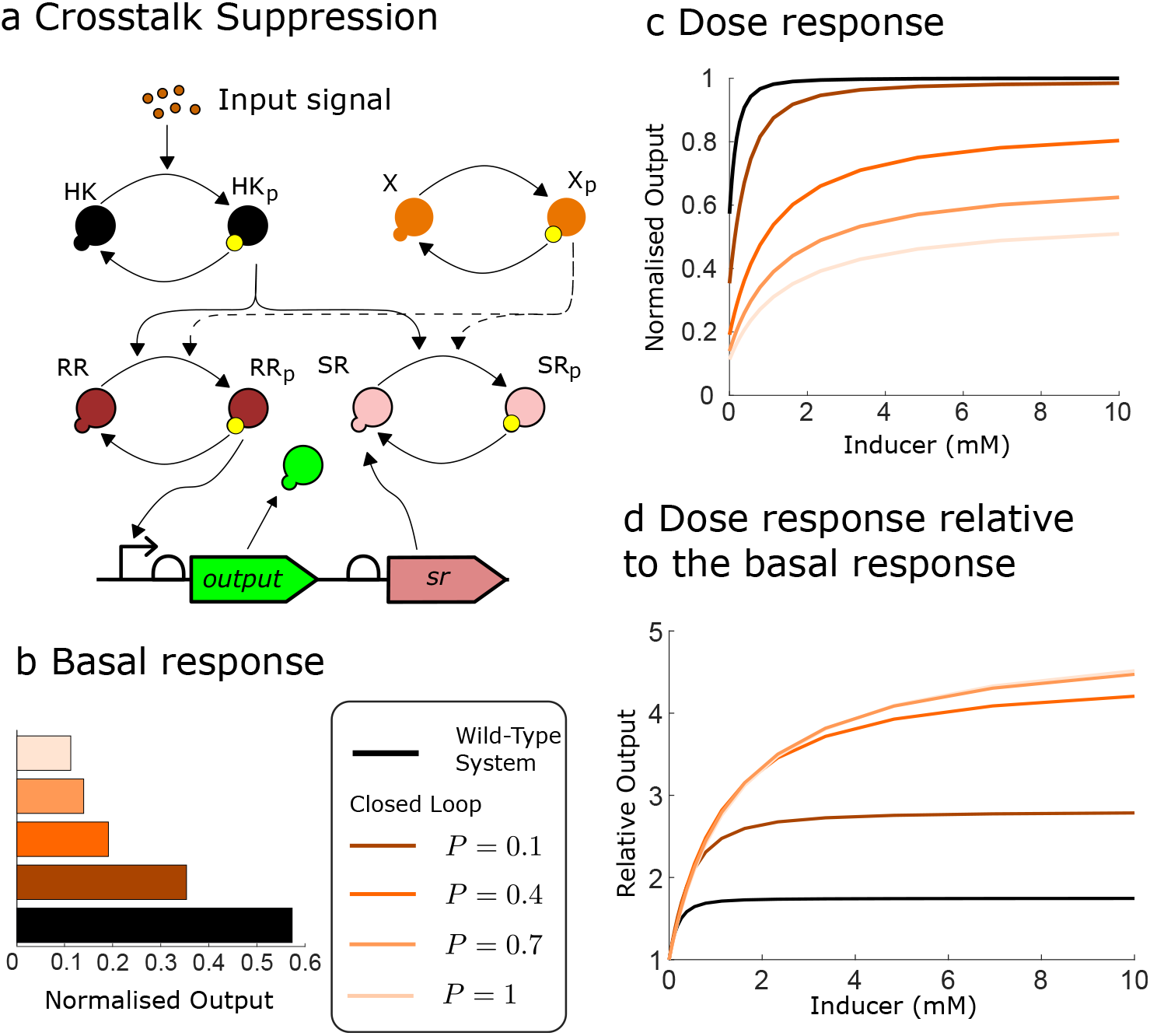
a. A schematic depiction of a two-component system with the dichotomous feedback architecture and crosstalk with another histidine kinase *X*. b. Basal response ([I] = 0 [mM]) of the wild-type and the closed-loop systems with various feedback strengths divided by the maximal response. c. Dose response curves of the wild-type and the closed-loop systems with various feedback strengths divided by the maximal response. d. Dose response curves of the wild-type and the closed-loop systems with various feedback strengths each divided by their basal response. The curves for *P* = 1 and *P* = 0.7 are indistinguishable by the naked eye. These simulations suggest that dichotomous feedback architecture can reduce crosstalk in the system.

## Discussion

In this paper we proposed a new feedback architecture realised using a signal sequestration motif, which is conceptually and architecturally different from previously proposed feedback mechanisms. We term this feedback mechanism *dichotomous feedback*. Sequestration motifs occur naturally in many biological systems and new ones can be designed by taking advantage of existing reactions in the process; this makes our dichotomous feedback practical to use. Our theoretical analysis was performed for two-component signalling systems composed of a response regulator and a histidine kinase with phosphatase abilities. In particular, we considered dephosphorylation enhancement, that can be realised by introducing another phosphatase, and phosphorylation sequestration, that can be realised by introducing another response regulator sequestering activating kinase.

Dephosphorylation enhancement was used in the past to build feedback in two-component systems and constitutes an example of dichotomous feedback based on the signal sequestration motif [Chang et al., 2013, Jones et al., 2021]. Understanding and studying the signal sequestration mechanism allowed us to propose a ‘dual’ dichotomous feedback architecture based on phosphorylation sequestration. We showed that phosphorylation sequestration and dephosphorylation enhancement architectures exhibit a qualitatively similar behaviour both in the absence and the presence of feedback. The difference between the two architectures lies in their effect on a particular system and either architecture can provide stronger feedback depending on the controlled system. The phosphorylation sequestration architecture appears to be more flexible as some response regulators (such as CheY_1_ in *S. meliloti*) may not have a phosphatase.

When binding between the phosphorylated histidine kinase and the second response regulator occurs, it renders the kinase inactive (until it is phosphorylated again). Sequestration – unlike annihilation – of the kinase gives more flexibility in the design of synthetic circuits, and troubleshooting can prove potentially simpler provided that there are means to measure the sequestered signal. Furthermore, annihilation and downregulation of histidine kinase molecules can have negative effects on the robustness of the two-component system as we demonstrated theoretically. The reason for such a behaviour appears to be architectural as annihilation of *only* phosphorylated kinases or phosphorylated response regulators exhibits similar robustness properties to dichotomous feedback. However, engineering a molecule targeting only the active version of a protein for the molecular sequestration based feedback may be more challenging than engineering a second response regulator for the dichotomous feedback.

As dichotomous feedback sequesters existing system fluxes, it also sequesters possible cross-talk fluxes affecting the system. Due to sequestration of cross-talk fluxes, dichotomous feedback can be designed to reduce the effect of cross-talk on the system. The design of a cross-talk reducing feedback is more delicate and was studied previously in [Steel et al., 2018, Steel et al., 2019]. In this work, we provided a signal sequestration interpretation of these results.

We focused on a two-component system in *E. coli*, however, our theoretical results can be applied to systems that share this architectural principle, such as sharing transcription and translation initiation molecules. Therefore, dichotomous feedback can potentially be realised in a wide variety of systems, which is of value if engineering molecular sequestration based feedback or transcription downregulation is not straightforward. We finally note that signal sequestration motifs can be used in other applications, for example, for tuning transcription initiation as was suggested in [Potvin-Trottier et al., 2016]. Furthermore, as two-component systems are found in all domains of life it would be interesting to explore the sequestration mechanism and dichotomous feedback in different cell types. The design would have to be adapted to the specificities of the system considered (for example, higher organisms often display multi-step phosphorelay architectures, which could also offer more tunability options to the design) but the overall architecture would hold. Examples include: osmolarity sensing in yeast, red/far-red light-sensing phytochromes in fungi and cytokinin signalling in plants [Wuichet et al.,2010, Schaller et al., 2011]. In these systems, a comprehensive study of the extrinsic noise properties of the dichotomous feedback mechanism would be needed, rather than just intrinsic noise as done in this paper. Metabolic pathways are the next obvious candidates for such feedback architectures to be implemented [Chubukov et al., 2012].

## Materials and Methods

### Mathematical Modelling

We used mass-action and Hill kinetic formalisms in order to model the chemical reactions. Noise analysis was performed using the Linear Noise Approximation of the Chemical Master Equation [Van Kampen, 1992] (see Section A.4.2 in SI) and the direct Gillespie algorithm for simulating the Chemical Master Equation implemented in [Zhou et al., 2011]. The robustness analysis was performed using the control-theoretic tool — the sensitivity function [Astrom and Murray, 2010] as described in Section A.3 in SI. The numerical computations were performed in MATLAB using a built-in ordinary differential equation solver ode15s. Code to produce all figures in the main text and the supplementary information can be found at https://github.com/oxfordcontrol/dichotomous-feedback.

## Author Contributions

ND and AS contributed equally to this work. Conceptualization: AS, ND, GHW and AP; Data Curation: AS, ND; Formal Analysis, AS, EA, HS, AP; Investigation: AS, ND, EA, AN, HS, AP; Methodology: AS, ND, EA, HS, GHW, AP; Writing - Original Draft: AS, AP; Writing - Review & Editing, AS, ND, EA, AN, HS, GHW, AP; Funding Acquisition, AP; Supervision: GHW, AP; Validation: EA, AP; Resources: GHW and AP. Correspondence should be addressed to Prof Papachristodoulou at.

## Acknowledgements

The authors would like to thank Dr M. Roberts for discussions regarding two-component systems. This work is supported by UK’s Engineering and Physical Sciences (EPSRC) Grant EP/M002454/1.

