## supplementary information for "Dichotomous Feedback: A Signal Sequestration-based Feedback Mechanism for Biocontroller Design"

**A Mathematical Preliminaries** 1

In Section A we introduce some mathematical concepts and analysis tools used throughout this SI. In Section B we model the phosphorylation sequestration motif and the closed loop system, which we refer to as SR dichotomous feedback. In Section C, we model the de-phosphorylation enhancement motif and the closed loop system, which we refer to as PH dichotomous feedback. In Section D we compare the feedbacks and perform additional simulations. The rest of the sections are devoted to technical derivations and information that support the main text.

### A Mathematical Preliminaries

#### A.1 Dynamical Systems

We consider a system in the following form

$$\dot{x} = f(x, u), \quad x(0) = x_0, \tag{S1a}$$

$$y = h(x) \tag{S1b}$$

where $f : \mathcal{D} \times \mathcal{U} \to \mathbb{R}^n$, $h : \mathbb{R}^n \to \mathbb{R}^m$ $u : \mathbb{R}_{\geq 0} \to \mathcal{U}$, $u$ belongs to the space $\mathcal{U}_\infty$ of Lebesgue measurable functions on $\mathcal{U}$, and $\mathcal{D}, \mathcal{U}$ are closed, convex subsets of $\mathbb{R}^n$ and $\mathbb{R}^k$, respectively. We denote the solutions to the system (S1a) with an initial condition $x_0$, and a control signal $u$ as $\phi(t, x_0, u)$ (mathematically, $\phi : \mathbb{R} \times \mathcal{D} \times \mathcal{U}_\infty \to \mathbb{R}^n$). We assume that $f \in C^2$ in both $u$ and $x$ on an open set containing $\mathcal{D} \times \mathcal{U}$. While our assumptions are quite restrictive, in general, they are fulfilled in our models. We call a set $\mathcal{S} \subseteq \mathcal{D}$ forward-invariant if $\phi(t, x, 0) \in \mathcal{S}$ for all $x \in \mathcal{S}$, and $t \geq 0$.

#### A.2 Monotone Systems Theory

In general, it is difficult to compare the trajectories $\phi(t, x^0)$ of dynamical systems knowing only the initial conditions $x^0$. However, for a monotone system the following simple relation holds: Let $z_i$ denote the $i$-th component of the vector $z$. If the system is monotone then if $x_i^0$ is larger than $x_i^1$ for all $i$ then $\phi_i(t, x^0)$ is larger than $\phi_i(t, x_i^1)$ for all $t > 0$. This property leads to a number of powerful results, which we elaborate in the sequel.

We will consider partial orders induced by orthants $\mathcal{K} = T_\mathcal{K}\mathbb{R}^n_{\geq 0}$, where $T_\mathcal{K} = \mathrm{diag}[(-1)^{\varepsilon_1}, \cdots, (-1)^{\varepsilon_n}]$, $\varepsilon_i \in \{0, 1\}$ and $\mathbb{R}^n_{\geq 0} = \{x \in \mathbb{R}^n | x_i \geq 0, \forall i = 1, \ldots, n\}$. The partial order on $\mathcal{K}$ is defined as $x \succeq_\mathcal{K} y$ if and only if $x - y \in \mathcal{K}$. Since in our case $\mathcal{K}$ is an orthant, the partial order is an element-wise comparison, that is, $x_i \geq y_i$ if $\varepsilon_i = 0$ and $x_i \leq y_i$ if $\varepsilon_i = 1$. The partial order on the space of signals $u \in \mathcal{U}_\infty$ is defined as $u_1 \preceq_\mathcal{K} u_2$ if $u_1(t) \preceq_\mathcal{K} u_2(t)$ holds for all $t \geq 0$. The set $[x,\ y]_\mathcal{K} = \{z \in \mathbb{R}^n | x \preceq_\mathcal{K} z \preceq_\mathcal{K} y\}$ is called *an order-interval.* Monotonicity for control systems is understood as preservation of partial orders in the initial conditions and inputs.

**Definition 1** *The system $\dot{x} = f(x, u)$ is called* monotone *on $\mathcal{X}, \mathcal{U}$ with respect to $\mathcal{K}_x$, $\mathcal{K}_u$ if $\phi(t, x, u) \preceq_{\mathcal{K}_x} \phi(t, y, v)$ for all $t \geq 0$, and for all $x \preceq_{\mathcal{K}_x} y$, and $u \preceq_{\mathcal{K}_u} v$, where $x,\ y \in \mathcal{X}$ and $u,\ v \in \mathcal{U}_\infty$.*

We now define *static characteristics* in the context of monotone control systems.

**Definition 2** *We say that a controlled dynamical system $\dot{x} = f(x, u)$ is endowed with the static Input/State characteristic $k_x(\cdot) : \mathcal{U} \to \mathcal{X}$ if for each constant input $u(t) \equiv \overline{u}$ there exists a (necessarily unique) globally asymptotically stable equilibrium $k_x(\overline{u})$. For systems with an output map $y = h(x)$, we also define the static Input/Output characteristic as $k_y(\overline{u}) = h(k_x(\overline{u}))$, provided that an Input/State characteristic exists and that $h$ is continuous.*

The systems preserving the partial order were fully characterised in [1] and the set for orthant-monotonicity becomes a simple check of the sign-pattern of the Jacobian.

**Theorem S1 ( [1])** *Consider the system $\dot{x} = f(x, u)$ under the assumptions above. The system is monotone with respect to* $\mathrm{diag}[(-1)^{\varepsilon_1}, \cdots, (-1)^{\varepsilon_n}]\mathbb{R}^n_{\geq 0}$, $\mathrm{diag}[(-1)^{\delta_1}, \cdots, (-1)^{\delta_k}]\mathbb{R}^k_{\geq 0}$, *where $\varepsilon_i$, $\delta_i$ are equal to zero or one, if and only if*

$$(-1)^{\varepsilon_i + \varepsilon_j} \frac{\partial f_i}{\partial x_j} \geq 0, \quad i \neq j \quad \forall (x, u) \in \mathrm{int}\{\mathcal{X}\} \times \mathrm{int}\{\mathcal{U}\},$$

$$(-1)^{\varepsilon_i + \delta_j} \frac{\partial f_i}{\partial u_j} \geq 0, \quad \forall (x, u) \in \mathcal{X} \times \mathrm{int}\{\mathcal{U}\},$$

*where $i,\ j = 1, \ldots, n$.*

Unforced monotone systems (i.e., $u = 0$) possess a number of strong stability properties [2]. For example:

- Chaotic behaviour is not possible in $\mathcal{X}$,
- There are no stable limit cycles in $\mathcal{X}$,
- If $x^*$ and $x^\bullet$ are asymptotically stable equilibria of the system and $x^* \ll_\mathcal{K} x^\bullet$, then for any $x$ such that $x \in [x^*, x^\bullet]_\mathcal{K}$ the flow $\phi(t, x)$ converges to $x^*$, $x^\bullet$ or there is an asymptotically stable equilibrium $x^\dagger$ such that $x^\dagger \in [x^*, x^\bullet]_\mathcal{K}$.

- Basins of attraction are unions of order-intervals [3].

Showing attractivity of monotone systems on an order-interval can be reduced to the computation of two trajectories. The argument is the following: let $x \preceq_{\mathcal{K}} y$, if $\phi(t, x)$, $\phi(t, y)$ converge to an equilibrium $x^*$, then the trajectories with initial conditions in $[x, y]_{\mathcal{K}}$ converge to $x^*$ by definition of monotonicity. If $x^*$ is additionally stable, then the order-interval $[x,\ y]_{\mathcal{K}}$ lies in the basin of attraction of an asymptotically stable equilibrium $x^*$ [4]. Further, using these two trajectories we can actually build a Lyapunov function as discussed in [5, 6].

A similar idea is used in the derivation of the small-gain result for monotone systems, that is, verification of attractivity of two monotone asymptotically stable systems. Consider the systems:

$$\dot{x}_1 = f_1(x_1, u_1),\ h_1(x_1) = y_1 = u_2 \tag{S2}$$

$$\dot{x}_2 = f_2(x_2, u_2),\ h_2(x_2) = y_2 = u_1 \tag{S3}$$

where $\mathcal{X}_1 \subseteq \mathbb{R}^{n_1}$, $\mathcal{U}_1 \subseteq \mathbb{R}^m$, $\mathcal{Y}_1 \subseteq \mathbb{R}^p$, $\mathcal{X}_2 \subseteq \mathbb{R}^{n_2}$, $\mathcal{U}_2 \subseteq \mathbb{R}^p$, $\mathcal{Y}_2 \subseteq \mathbb{R}^m$, and $\mathcal{Y}_2 \subseteq \mathcal{U}_1$, $\mathcal{Y}_1 \subseteq \mathcal{U}_2$. We assume that both systems are input-output monotone with respect to the orthants and $\mathcal{X}_i$, $\mathcal{U}_i$, $\mathcal{Y}_i$ are Cartesian products of (semi-) closed intervals (for simplicity, we will call them boxes). The orthants for the state, control and output orders are given as $\mathcal{K}_{\mathcal{X}_1}$, $\mathcal{K}_{\mathcal{U}_1}$, $\mathcal{K}_{\mathcal{Y}_1}$ for the system (S2) and $\mathcal{K}_{\mathcal{X}_2}$, $\mathcal{K}_{\mathcal{U}_2}$, $\mathcal{K}_{\mathcal{Y}_2}$ for the system (S3). Let $T_1$ and $T_2$ be diagonal matrices with entries from $\{1, -1\}$ such that $\mathcal{K}_{\mathcal{U}_1} = T_1 \mathcal{K}_{\mathcal{Y}_2}$ and $\mathcal{K}_{\mathcal{U}_2} = T_2 \mathcal{K}_{\mathcal{Y}_1}$. Let also $T_i^+ = \max\{T_i, 0\}$ and $T_i^- = -\min\{T_i, 0\}$, where the minimum and the maximum are interpreted entry-wise, and consider a function $\eta_1 : \mathbb{R}^{2p} \to \mathbb{R}^{2p}$ such that $\eta_1(a, b) = \begin{pmatrix}(T_1^+ a + T_1^- b)^T & (T_1^+ b + T_1^- a)^T\end{pmatrix}^T$. We define the function $\eta_2 : \mathbb{R}^{2m} \to \mathbb{R}^{2m}$ using $T_2$ in an analogous manner. Now we can formulate the monotone system small-gain theorem:

**Theorem S2 ( [7])** *Consider a well-posed interconnection of I/O orthant-monotone systems (S2,S3), and assume every solution* $\begin{pmatrix}x_1^T & x_2^T\end{pmatrix}^T$ *is bounded. Suppose both I/O systems admit continuous steady-state I/O characteristics* $\gamma_1 : \mathcal{U}_1 \to \mathcal{Y}_1$, $\gamma_2 : \mathcal{U}_2 \to \mathcal{Y}_2$ *as well as continuous input-state characteristics. Define the discrete iteration that given* $u_1^-(0) \preceq_{\mathcal{K}_{\mathcal{U}_1}} u_1^+(0)$ *in* $\mathcal{U}_1$ *calculates input and output values for every* $k = 0, 1, 2...$ *as follows:*

$$\begin{aligned}
&y_1^-(x) = \gamma^1(u_1^-(k)), y_1^+(x) = \gamma^1(u_1^+(k)),\\
&\begin{pmatrix} u_2^-(k) \\ (u_2^+(k) \end{pmatrix} = \eta_1(y_1^-(k), y_1^+(k))\\
&y_2^-(x) = \gamma^2(u_2^-(k)), y_2^+(x) = \gamma^2(u_2^+(k)),\\
&\begin{pmatrix} u_1^-(k+1) \\ u_1^+(k+1) \end{pmatrix} = \eta_2(y_2^-(k), y_2^+(k)),
\end{aligned} \tag{S4}$$

*where* $\eta_1$, $\eta_2$ *are defined above. Provided that for all initial conditions* $u_1^-(0)$, $u_1^+(0)$ *the discrete iteration converges towards a unique equilibrium* $\begin{pmatrix}\overline{u}^T & \overline{u}^T\end{pmatrix}^T$, *the closed loop system is globally convergent. Namely, for every solution* $\begin{pmatrix}x_1^T & x_2^T\end{pmatrix}^T$ *of (S2,S3), the functions* $y_i = h_i(x_i(t))$ *fulfill*

$$\lim_{t\to\infty} \begin{pmatrix}y_1^T & y_2^T\end{pmatrix}^T = \begin{pmatrix}\gamma_1(\overline{u})^T & \overline{u}^T\end{pmatrix}^T.$$

*Also, every state solution* $\begin{pmatrix}x_1^T & x_2^T\end{pmatrix}^T$ *converges towards a unique globally attractive equilibrium.*

When the feedback interconnection is static, that is $y_2 = u_2$, the result can be relaxed by using the following discrete iteration

$$\begin{aligned}
&y_1^-(x) = \gamma^1(u_1^-(k)), y_1^+(x) = \gamma^1(u_1^+(k)),\\
&\begin{pmatrix}(u_1^-(k+1))^T & (u_1^+(k+1))^T\end{pmatrix}^T = \eta(y_1^-(k), y_1^+(k)),
\end{aligned} \tag{S5}$$

where $\eta$ is defined using $T$ such that $\mathcal{K}_{\mathcal{U}_1} = T\mathcal{K}_{\mathcal{Y}_1}$ (see [7]). Furthermore, if the system is additionally single-input-single-output, that is $\dot{x} = f(x, u)$, $h(x) = y = u : \mathcal{X} \to \mathbb{R}$, then we can drop the requirement on $\mathcal{X}$ being rectangular, which allows to apply the results to wider class of systems (see [1] or Proposition 1 in [7]).

#### A.3 Transfer Functions

We study deviations of the model (S1) from the steady-state $x^*$ taken for a particular constant value of $u(t)$ denoted as $u^*$ (i.e., we assume $u(t) = u^*$ for all $t \geq 0$). In particular, we linearise the model about $(x^*, u^*)$ as follows:

$$\frac{d}{dt}\Delta x = A\Delta x + B\Delta u,$$
$$\Delta y = C\Delta x,$$

where the matrices $A = \left.\frac{\partial f(x,u)}{\partial x}\right|_{(x,u)=(x^*,u^*)}$, $B = \left.\frac{\partial f(x,u)}{\partial u}\right|_{(x,u)=(x^*,u^*)}$, and $C = \left.\frac{dh(x)}{dx}\right|_{x=x^*}$ are the first terms of the Taylor expansion of $f(x,u)$ and $h(x)$ about $(x^*, u^*)$. If the vector-valued functions $f(x,u)$, $h(x)$ are sufficiently smooth and the matrix $A$ does not have eigenvalues on the imaginary axis, then this linearisation describes the behaviour of the nonlinear model locally around the steady-state $(x^*, u^*)$.

In control theory, taking the Laplace transform of the linear model often facilitates analysis. In particular, we obtain *the transfer function* $G(s)$, which describes the linear relation between the inputs $\Delta U(s)$, which is a signal $\Delta u(t)$ after the Laplace transformation, and the outputs $\Delta Y(s)$, which is a signal $\Delta y(t)$ after the Laplace transformation:

$$\Delta Y(s) = \underbrace{C(sI - A)^{-1}B}_{G(s)}\Delta U(s).$$

The magnitude of the transfer function at $s = \jmath\omega$ (where $\jmath$ is the complex identity and $\omega$ is a nonnegative and real number) describes how the system reacts to sinusoidal signals of frequency $\omega$, e.g., $\sin(\omega t)$. If the magnitude $\|G(\jmath\omega)\|_2$ (computed as the maximal singular value of $G(\jmath\omega)$) is larger than one then this signal is amplified and when the magnitude is lower than one then the signal is attenuated.

Transfer functions are a useful tool for evaluating the robustness of the closed loop system around the steady-state. In this setting, we assume that the output $y$ of the controlled system called process $G$ is used by the controller $K$ in order to compute the signal $u$, which modifies the behaviour of the process $G$. This creates a feedback loop, which we can study using transfer functions. In particular, consider the setting in Figure S1, which we use in this paper. The inputs and outputs are related using the following equation

$$Y(s) = \underbrace{(I - G_{\mathrm{yu}}(s)K(s))^{-1}G_{yz}(s)}_{T(s)}Z(s) + \underbrace{(I - G_{\mathrm{yu}}(s)K(s))^{-1}}_{S(s)}W(s),$$

where $G_{\mathrm{yu}}$, $G_{\mathrm{yz}}$ are the transfer functions from $U(s)$ to $Y(s)$ and $Z(s)$ to $Y(s)$, respectively; $W(s)$ models the process disturbances (such as, modelling errors), $Z(s)$ models the external inputs and $Y(s)$ models the output of the process.

The function $S(s)$ describes the contribution of the disturbance $W(s)$ to the output, which ideally should either attenuate the disturbances (i.e., the magnitude of $S(\jmath\omega)$ for $\omega \in \mathbb{R}$ is close to zero) or at least not amplify them (i.e., the magnitude of $S(\jmath\omega)$ for $\omega \in \mathbb{R}$ is close to one). In control engineering, the rule-of-thumb is to design the controller $K$ so that $\|S(s)\|_{\mathcal{H}_\infty} = \max_{\omega\in\mathbb{R}} \|S(\jmath\omega)\|_2 \leq 1.2$ [8]. Furthermore, for low frequencies the magnitude of $S(s)$ should be as small as possible and smaller than one.

The function $T(s)$ describes the behaviour of the system with respect to the external inputs $Z(s)$. In our case, the external inputs can be the concentration of the inducer and the production rate of certain proteins.

#### A.4 Chemical Reaction Modelling

##### A.4.1 Deterministic Modelling

Consider the biochemical species $X_1, \ldots, X_n$ with concentrations $x_1, \ldots, x_n$ participating in the following chemical reactions:

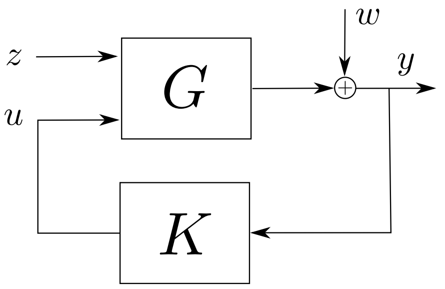

$$S = (I - G_{yu}K)^{-1}$$
$$T = (I - G_{yu}K)^{-1}G_{yz}$$

Figure S1: Block diagram of the process $G$ to be controlled and the controller $K$ feedback loop used in this paper. The signal $z$ signifies the external inputs, $y$ the measured output, $w$ are the process disturbances (e.g., modelling errors or external perturbations) and $u$ is the control signal computed by the controller $K$. The significance of the functions $S$ and $T$ is discussed in the text below.

$$R_j: \quad \sum_{i=1}^{n} \alpha_{ij} X_i \xrightarrow{v_j} \sum_{i=1}^{n} \beta_{ij} X_i \quad j = 1, \ldots, m$$

where $\alpha_{ij}$, $\beta_{ij}$ are nonnegative integers and $v_j$ are the reactions rates. We assume that there are at most two reactants and two products in every reaction, hence for every $j$ there are at most two nonzero $\alpha_{ij}$ and $\beta_{ij}$. In the case of mass-action modelling, the reaction rates $v_j$ depend on the concentration $x_i$ for $i$ with nonzero $\alpha_{ij}$ as follows:

$$v_j(x) = k_{ij} \prod_{i=1}^{n} x_i^{\alpha_{ij}}.$$

If Michaelis-Menten or Hill kinetics are assumed then $v_j(x)$ is a rational function of particular concentrations $x_i$.

The standard deterministic reaction model can be written as follows:

$$\dot{x} = Sv(x), \tag{S6}$$

where $x$ is the vector of concentrations, $v(x)$ is the vector-function of propensities with entries $v_j$ and $S$ is called the stoichiometric matrix defined as:

$$S_{ij} = \beta_{ij} - \alpha_{ij}.$$

##### A.4.2 Linear Noise Approximation

We evaluate the stochastic properties of the system using the Linear Noise Approximation of the Chemical Master Equation. If we consider the deterministic model (S6) with the stoichiometric matrix $S$ and the propensity vector $v(x)$, the linear noise approximation is given by:

$$\dot{\xi} = A(x)\xi + B(x)\dot{w}, \tag{S7}$$

where $A(x)$ is the Jacobian of $Sv(x)$, $w$ is the Gaussian white noise, $B(x) = S(\mathrm{diag}\{v(x)\})^{1/2}$, where $\mathrm{diag}\{z\}$ is the matrix with the elements of the vector $z$ on the diagonal and the operator $\cdot^{1/2}$ is the element-wise square root. The dynamics of the noise vector $\xi$ (S7) are linear with the zero mean and covariance $P = \mathbb{E}(\xi\xi^T)$, which can be computed using the following matrix-valued differential equation

$$\dot{P} = PA^T(x(t)) + A(x(t))P + B(x(t))B^T(x(t)), \tag{S8}$$

where the trajectory $x(t)$ obeys (S6). In stationarity the covariance matrix can be computed using standard linear algebra tools as it becomes the Lyapunov equation

$$PA^T(x^*) + A(x^*)P + B(x^*)B^T(x^*) = 0, \tag{S9}$$

where $x^*$ is the steady-state of (S6). The diagonal of the matrix $P$ contains the variance of particular species, while the off-diagonal elements signify covariance between different species.

### B Phosphorylation Sequestration Motif Modelling

#### B.1 Basic Model

We reproduce the model that was developed in the main text here for convenience. The chemical reaction model describing a two-component system can be written as follows:

$$\begin{aligned}
&\mathrm{HK} \xrightarrow{k_{\mathrm{ap}}(\mathrm{I})} \mathrm{HK_p}, \\
&\mathrm{HK_p} + \mathrm{RR} \xrightarrow{k_t} \mathrm{HK} + \mathrm{RR_p}, \qquad \mathrm{HK} + \mathrm{RR_p} \xrightarrow{k_p} \mathrm{HK} + \mathrm{RR}, \\
&\mathrm{RR},\ \mathrm{RR_p},\ \mathrm{Output},\ \mathrm{HK},\ \mathrm{HK_p} \xrightarrow{\delta} \emptyset, \quad \emptyset \xrightarrow{\beta_{\mathrm{RR}}} \mathrm{RR},\ \emptyset \xrightarrow{\beta_{\mathrm{HK}}} \mathrm{HK},\ \emptyset \xrightarrow{k_{\mathrm{out}}([\mathrm{RR_p}])} \mathrm{Output},
\end{aligned} \tag{S10}$$

where the suffix $\cdot_{\mathrm{p}}$ denotes the phosphorylated form of a protein. We assumed that the autophosphorylation rate $k_{\mathrm{ap}}$ depends on the inducer concentration as a Michaelis-Menten function, while the output expression initiation depends on the response regulator as a Hill function, as previously suggested [9]:

$$\begin{aligned}
k_{\mathrm{ap}}(\mathrm{I}) &= k_{\mathrm{ap-max}} \frac{[\mathrm{I}]}{[\mathrm{I}] + K_{\mathrm{da}}}, \\
k_{\mathrm{out}}([\mathrm{RR_p}]) &= k_{\mathrm{out-max}} \frac{([\mathrm{RR_p}]/K_{\mathrm{dr}})^n}{([\mathrm{RR_p}]/K_{\mathrm{dr}})^n + 1},
\end{aligned}$$

where $k_{\mathrm{ap-max}}$ is the maximal autophosphorylation rate, $K_{\mathrm{da}}$ is the inducer dissociation constant, $k_{\mathrm{out-max}}$ is the maximum production rate of the output protein, $K_{\mathrm{dr}}$ is the dissociation constant, and $n$ is the cooperativity coefficient. We modelled the reactions for the protein SR using the following chemical reactions:

$$\begin{aligned}
&\mathrm{HK_p} + \mathrm{SR} \xrightarrow{k_{tc}} \mathrm{HK} + \mathrm{SR_p}, \quad \mathrm{HK} + \mathrm{SR_p} \xrightarrow{k_{pc}} \mathrm{HK} + \mathrm{SR}, \\
&\mathrm{SR}, \mathrm{SR_p} \xrightarrow{\delta} \emptyset, \qquad \emptyset \xrightarrow{\beta_{\mathrm{SR}}} \mathrm{SR}.
\end{aligned} \tag{S11}$$

Assuming these chemical reactions and mass-action kinetics, we can obtain the following differential equation model:

$$\begin{aligned}
\frac{d[\mathrm{HK}]}{dt} &= \beta_{\mathrm{HK}} - \delta[\mathrm{HK}] - k_{\mathrm{ap}}[\mathrm{HK}] + k_t[\mathrm{HK_p}][\mathrm{RR}] + k_{tc}[\mathrm{HK_p}][\mathrm{SR}], \\
\frac{d[\mathrm{HK_p}]}{dt} &= -k_t[\mathrm{HK_p}][\mathrm{RR}] - k_{tc}[\mathrm{HK_p}][\mathrm{SR}] + k_{\mathrm{ap}}[\mathrm{HK}] - \delta[\mathrm{HK_p}], \\
\frac{d[\mathrm{RR}]}{dt} &= \beta_{\mathrm{RR}} - \delta[\mathrm{RR}] - k_t[\mathrm{HK_p}][\mathrm{RR}] + k_p[\mathrm{HK}][\mathrm{RR_p}], \\
\frac{d[\mathrm{RR_p}]}{dt} &= -\delta[\mathrm{RR_p}] + k_t[\mathrm{HK_p}][\mathrm{RR}] - k_p[\mathrm{HK}][\mathrm{RR_p}], \\
\frac{d[\mathrm{SR}]}{dt} &= \beta_{\mathrm{SR}} - \delta[\mathrm{SR}] - k_{tc}[\mathrm{HK_p}][\mathrm{SR}] + k_{pc}[\mathrm{HK}][\mathrm{SR_p}], \\
\frac{d[\mathrm{SR_p}]}{dt} &= -\delta[\mathrm{SR_p}] + k_{tc}[\mathrm{HK_p}][\mathrm{SR}] - k_{pc}[\mathrm{HK}][\mathrm{SR_p}], \\
\frac{d[\mathrm{Output}]}{dt} &= k_{\mathrm{out}}([\mathrm{RR_p}]) - \delta[\mathrm{Output}].
\end{aligned} \tag{S12}$$

#### B.2 Model Simplification

We first consider the case in which the production rate of the protein SR is constant and equal to $\beta_{\text{SR}}$, as in the model above. In the sequestration mechanism model, we omit the output protein, since we want to study the dynamics of the phosphorylated response regulator. We proceed by introducing the following variables:

$$\begin{aligned}[\text{HK}_{\text{sum}}] &= [\text{HK}] + [\text{HK}_{\text{p}}],\\ [\text{RR}_{\text{sum}}] &= [\text{RR}] + [\text{RR}_{\text{p}}],\\ [\text{SR}_{\text{sum}}] &= [\text{SR}] + [\text{SR}_{\text{p}}],\end{aligned}$$

which can be used to simplify the model. These obey the following differential equations:

$$\begin{aligned}\frac{d[\text{HK}_{\text{sum}}]}{dt} &= \beta_{\text{HK}} - \delta[\text{HK}_{\text{sum}}],\\ \frac{d[\text{RR}_{\text{sum}}]}{dt} &= \beta_{\text{RR}} - \delta[\text{RR}_{\text{sum}}],\\ \frac{d[\text{SR}_{\text{sum}}]}{dt} &= \beta_{\text{SR}} - \delta[\text{SR}_{\text{sum}}],\end{aligned}$$

where in the steady-state we have $[\text{HK}_{\text{sum}}]^* = \text{HK}_{\text{tot}} = \beta_{\text{HK}}/\delta$, $[\text{RR}_{\text{sum}}]^* = \text{RR}_{\text{tot}} = \beta_{\text{RR}}/\delta$, $[\text{SR}_{\text{sum}}]^* = \text{SR}_{\text{tot}} = \beta_{\text{SR}}/\delta$. This implies that *in the steady-state*, we have the conservation laws:

$$[\text{HK}]^* + [\text{HK}_{\text{p}}]^* = \text{HK}_{\text{tot}}, \tag{S13}$$

$$[\text{RR}]^* + [\text{RR}_{\text{p}}]^* = \text{RR}_{\text{tot}}, \tag{S14}$$

$$[\text{SR}]^* + [\text{SR}_{\text{p}}]^* = \text{SR}_{\text{tot}}. \tag{S15}$$

The introduction of the new variables allows us to eliminate the variables $[\text{HK}_{\text{p}}]$, $[\text{RR}_{\text{p}}]$ and $[\text{SR}_{\text{p}}]$ from the system of equations, while adding the differential equations for the total concentrations of the proteins. In control-theoretic terms, this operation is simply a *state-space transformation* that results in an equivalent set of equations. This gives the following simplified model:

$$\begin{aligned}\frac{d[\text{HK}]}{dt} &= \beta_{\text{HK}} - \delta[\text{HK}] - k_{\text{ap}}[\text{HK}] + k_t([\text{HK}_{\text{sum}}] - [\text{HK}])[\text{RR}] + k_{tc}([\text{HK}_{\text{sum}}] - [\text{HK}])[\text{SR}],\\ \frac{d[\text{RR}]}{dt} &= \beta_{\text{RR}} - \delta[\text{RR}] - k_t([\text{HK}_{\text{sum}}] - [\text{HK}])[\text{RR}] + k_p[\text{HK}]([\text{RR}_{\text{sum}}] - [\text{RR}]),\\ \frac{d[\text{SR}]}{dt} &= \beta_{\text{SR}} - \delta[\text{SR}] - k_{tc}([\text{HK}_{\text{sum}}] - [\text{HK}])[\text{SR}] + k_{pc}[\text{HK}]([\text{SR}_{\text{sum}}] - [\text{SR}]),\\ \frac{d[\text{HK}_{\text{sum}}]}{dt} &= \beta_{\text{HK}} - \delta[\text{HK}_{\text{sum}}],\\ \frac{d[\text{RR}_{\text{sum}}]}{dt} &= \beta_{\text{RR}} - \delta[\text{RR}_{\text{sum}}],\\ \frac{d[\text{SR}_{\text{sum}}]}{dt} &= \beta_{\text{SR}} - \delta[\text{SR}_{\text{sum}}].\end{aligned} \tag{S16}$$

Note that instead of eliminating $[\text{HK}_{\text{p}}]$, $[\text{RR}_{\text{p}}]$ and $[\text{SR}_{\text{p}}]$, we could have eliminated [HK], [RR] and [SR] or [HK], $[\text{RR}_{\text{p}}]$ and $[\text{SR}_{\text{p}}]$. In general, we could eliminate one version (phosphorylated or not) of histidine kinase, response regulator and sequestration protein. If $\beta_{\text{SR}}$ is not constant (e.g., the feedback is active), then we can still use the model (S16), however, the conservation law for the proteins $[\text{SR}] + [\text{SR}_{\text{p}}] = \text{SR}_{\text{tot}}$ will not be valid. In that case we have

$$\frac{d[\text{SR}_{\text{sum}}]}{dt} = Pk_{\text{out}}([\text{RR}_{\text{p}}] - \delta[\text{SR}_{\text{sum}}]). \tag{S17}$$

As a result, at the steady state: $[\text{SR}]^* + [\text{SR}_{\text{p}}]^* = [\text{SR}_{\text{sum}}] = Pk_{\text{out}}([\text{RR}_{\text{p}}]^*)/\delta$.

#### B.3 Properties of the Sequestration Motif and the Closed Loop Models.

This section largely follows similar developments in [10, 11], where monotone systems theory is used for systems analysis. First let us consider a simpler system:

$$\begin{aligned}
\frac{d[\mathrm{HK}]}{dt} &= \beta_{\mathrm{HK}} - \delta[\mathrm{HK}] - k_{\mathrm{ap}}[\mathrm{HK}] + k_t(\mathrm{HK}_{\mathrm{tot}} - [\mathrm{HK}])[\mathrm{RR}] + k_{tc}(\mathrm{HK}_{\mathrm{tot}} - [\mathrm{HK}])[\mathrm{SR}],\\
\frac{d[\mathrm{RR}]}{dt} &= \beta_{\mathrm{RR}} - \delta[\mathrm{RR}] - k_t(\mathrm{HK}_{\mathrm{tot}} - [\mathrm{HK}])[\mathrm{RR}] + k_p[\mathrm{HK}](\mathrm{RR}_{\mathrm{tot}} - [\mathrm{RR}]),\\
\frac{d[\mathrm{SR}]}{dt} &= \beta_{\mathrm{SR}} - \delta[\mathrm{SR}] - k_{tc}(\mathrm{HK}_{\mathrm{tot}} - [\mathrm{HK}])[\mathrm{SR}] + k_{pc}[\mathrm{HK}](\mathrm{SR}_{\mathrm{tot}} - [\mathrm{SR}]),
\end{aligned} \tag{S18}$$

where $\mathrm{HK}_{\mathrm{tot}} = \beta_{\mathrm{HK}}/\delta$, $\mathrm{RR}_{\mathrm{tot}} = \beta_{\mathrm{RR}}/\delta$ and $\mathrm{SR}_{\mathrm{tot}} = \beta_{\mathrm{SR}}/\delta$. We will the study its properties on the set $\mathcal{S}$:

$$\mathcal{S} = \left\{ \begin{pmatrix}[\mathrm{HK}] & [\mathrm{RR}] & [\mathrm{SR}]\end{pmatrix}^T \,\middle|\, 0 \le [\mathrm{HK}] \le \mathrm{HK}_{\mathrm{tot}}, 0 \le [\mathrm{RR}] \le \mathrm{RR}_{\mathrm{tot}}, 0 \le [\mathrm{SR}] \le \mathrm{SR}_{\mathrm{tot}} \right\}.$$

**Proposition S1** *Consider the phosporylation sequestration motif (i.e., system* (S18)*) with nonnegative parameters then*

1. *trajectories of the system* (S18) *cannot leave the set* $\mathcal{S}$*;*
2. *consider the natural two component system, i.e., the system* (7) *with* $\beta_{\mathrm{SR}} = 0$*. The equilibrium* $[\mathrm{HK}]^*$, $[\mathrm{RR}]^*$ *of this system is locally asymptotically stable if*

   $$(\delta + k_{\mathrm{ap}} + k_t[\mathrm{RR}]^*)(\delta + k_t[\mathrm{HK_p}]^* + k_p[\mathrm{HK}]^*) > k_t[\mathrm{HK_p}]^*(k_t[\mathrm{RR}]^* + k_p[\mathrm{RR_p}]^*),$$

   *where* $[\mathrm{HK_p}]^* = \mathrm{HK}_{\mathrm{tot}} - [\mathrm{HK}]^*$, $[\mathrm{RR_p}]^* = \mathrm{RR}_{\mathrm{tot}} - [\mathrm{RR}]^*$*. In particular, this condition is satisfied if:*

   $$1 > \frac{[\mathrm{HK_p}]^*}{\mathrm{HK}_{\mathrm{tot}}} + \frac{[\mathrm{RR_p}]^*}{\mathrm{RR}_{\mathrm{tot}}}.$$
3. *the system* (S18) *is not chaotic and cannot have stable limit cycles;*
4. *consider two trajectories of the system* (S18) $\phi_1(t, x_1)$ *with* $\phi_2(t, x_2)$ *with* $x_1 = \begin{pmatrix}[\mathrm{HK}] & [\mathrm{RR}] & [\mathrm{SR}]\end{pmatrix} = \begin{pmatrix}0 & 0 & 0\end{pmatrix}$ *and* $x_2 = \begin{pmatrix}[\mathrm{HK}] & [\mathrm{RR}] & [\mathrm{SR}]\end{pmatrix} = \begin{pmatrix}\mathrm{HK}_{\mathrm{tot}} & \mathrm{RR}_{\mathrm{tot}} & \mathrm{SR}_{\mathrm{tot}}\end{pmatrix}$*. If both these trajectories converge to the same point* $x^*$ *as* $t$ *grows to infinity, then* $x^*$ *is globally attractive in* $\mathcal{S}$*;*
5. *if the system* (S18) *has a globally asymptotically stable in* $\mathcal{S}$ *equilibrium for* $\beta_{\mathrm{SR}} \in \mathcal{P}$*, where* $\mathcal{P}$ *is an interval in* $\mathbb{R}_{\ge 0}$*, then the steady-state concentration of* [RR] *increases monotonically with* $\beta_{\mathrm{SR}} \in \mathcal{P}$*.*

*Proof:*

1. The first claim follows if the system is forward invariant on $\mathcal{S}$. We need to show that the vector field is point inwards on the boundary of $\mathcal{S}$. Since $\mathcal{S}$ is a box all we need to check is the sign of the functions $f_{\mathrm{HK}}$, $f_{\mathrm{RR}}$, $f_{\mathrm{SR}}$, where $d[\mathrm{HK}]/dt = f_{\mathrm{HK}}$ for $[\mathrm{HK}] = 0$ and $[\mathrm{HK}] = \mathrm{HK}_{\mathrm{tot}}$, $d[\mathrm{RR}]/dt = f_{\mathrm{RR}}$ for $[\mathrm{RR}] = 0$ and $[\mathrm{RR}] = \mathrm{RR}_{\mathrm{tot}}$, $d[\mathrm{SR}]/dt = f_{\mathrm{SR}}$ for $[\mathrm{SR}] = 0$ and $[\mathrm{SR}] = \mathrm{SR}_{\mathrm{tot}}$. The signs can be verified to be as follows:

   $$\begin{aligned}
   f_{\mathrm{HK}}\Big|_{[\mathrm{HK}]=0} &\ge 0, & f_{\mathrm{HK}}\Big|_{[\mathrm{HK}]=0} &\le 0\\
   f_{\mathrm{RR}}\Big|_{[\mathrm{RR}]=0} &\ge 0, & f_{\mathrm{RR}}\Big|_{[\mathrm{RR}]=\mathrm{RR}_{\mathrm{tot}}} &\le 0\\
   f_{\mathrm{SR}}\Big|_{[\mathrm{SR}]=0} &\ge 0, & f_{\mathrm{SR}}\Big|_{[\mathrm{SR}]=\mathrm{SR}_{\mathrm{tot}}} &\le 0
   \end{aligned}$$
2. Since the system becomes two dimensional with $\beta_{\mathrm{SR}} = 0$, it is locally asymptotically stable if the trace and determinant of the Jacobian at the steady state are negative We can compute the Jacobian as

   $$J = \begin{pmatrix} -J_{11} & -J_{12} \\ -J_{21} & -J_{22} \end{pmatrix},$$

where

$$J_{11} = \delta + k_{\text{ap}} + k_t[\text{RR}^*], \qquad J_{12} = k_t(\text{HK}_{\text{tot}} - [\text{HK}]^*),$$
$$J_{21} = k_t[\text{RR}]^* + k_p(\text{RR}_{\text{tot}} - [\text{RR}]^*), \qquad J_{22} = \delta + k_t(\text{HK}_{\text{tot}} - [\text{HK}]^*) + k_p[\text{HK}]^*,$$

and the results follows after computing the determinant since trace is negative.

As for the second part of the statement for nonnegative parameters and steady-states we have:

$$(\delta + k_{\text{ap}} + k_t(\text{RR}_{\text{tot}} - [\text{RR}_\text{p}]^*) + k_{tc}[\text{SR}]^*)(\delta + k_t(\text{HK}_{\text{tot}} - [\text{HK}]^*) + k_p[\text{HK}]^*) >$$
$$k_t(\text{RR}_{\text{tot}} - [\text{RR}_\text{p}]^*)(k_t(\text{HK}_{\text{tot}} - [\text{HK}]^*) + k_p[\text{HK}]^*).$$

Therefore, if

$$k_t(\text{RR}_{\text{tot}} - [\text{RR}_\text{p}]^*)(k_t(\text{HK}_{\text{tot}} - [\text{HK}]^*) + k_p[\text{HK}]^*) > k_t(\text{HK}_{\text{tot}} - [\text{HK}]^*)(k_t(\text{RR}_{\text{tot}} - [\text{RR}_\text{p}]^*) + k_p[\text{RR}_\text{p}]^*),$$

we also have stability. We have the following chain of equivalent identities:

$$k_p k_t(\text{RR}_{\text{tot}} - [\text{RR}_\text{p}]^*)[\text{HK}]^* > k_p k_t(\text{HK}_{\text{tot}} - [\text{HK}]^*)[\text{RR}_\text{p}]^*,$$
$$\Updownarrow$$
$$\text{RR}_{\text{tot}}[\text{HK}]^* - [\text{RR}_\text{p}]^*[\text{HK}]^* > \text{HK}_{\text{tot}}[\text{RR}_\text{p}]^* - [\text{HK}]^*[\text{RR}_\text{p}]^*$$
$$\Updownarrow$$
$$\frac{[\text{HK}]^*}{\text{HK}_{\text{tot}}} > \frac{[\text{RR}_\text{p}]^*}{\text{RR}_{\text{tot}}}$$
$$\Updownarrow$$
$$1 > \frac{[\text{HK}_\text{p}]^*}{\text{HK}_{\text{tot}}} + \frac{[\text{RR}_\text{p}]^*}{\text{RR}_{\text{tot}}}.$$

3.-4. These points follow directly from system's monotonicity on $\mathcal{S}$ with respect to the orthant $\mathbb{R}^3_{\geq 0}$. Let us show monotonicity using the Jacobian criterion. The Jacobian of the vector field in (S16) can be computed as follows

$$J_{11} = \delta + k_{\text{ap}} + k_t[\text{RR}] + k_{tc}[\text{SR}], \quad J_{12} = k_t(\text{HK}_{\text{tot}} - [\text{HK}]), \quad J_{13} = k_t(\text{HK}_{\text{tot}} - [\text{HK}]),$$
$$J_{21} = k_t[\text{RR}] + k_p(\text{RR}_{\text{tot}} - [\text{RR}]), \quad J_{22} = \delta + k_t(\text{HK}_{\text{tot}} - [\text{HK}]) + k_p[\text{HK}],$$
$$J_{31} = k_{tc}[\text{SR}] + k_{pc}(\text{SR}_{\text{tot}} - [\text{SR}]), \quad J_{33} = \delta + k_{tc}(\text{HK}_{\text{tot}} - [\text{HK}]) + k_{pc}[\text{HK}],$$

and we have

$$J = \begin{pmatrix} -J_{11} & J_{12} & J_{13} \\ J_{21} & -J_{22} & 0 \\ J_{31} & 0 & -J_{33}, \end{pmatrix}$$

where all $J_{ij}$ are nonnegative for all points in $\mathcal{S}$. Hence the system is monotone according to Theorem S1 in Section A.2.

5. The steady-state monotonicity follows from monotonicity with respect to the parameter $\beta_{\text{SR}}$. The gradient of the vector field with respect to $\beta_{\text{SR}}$ gives:

$$B = \begin{pmatrix} 0 \\ 0 \\ 1 + k_{pc}[\text{HK}]/\delta, \end{pmatrix}$$

and hence the system is control-monotone on $\mathbb{R}^3_{\geq 0} \times \mathbb{R}_{\geq 0}$ according to Theorem S1 in Section A.2.

■

In order to determine close-loop stability for a specific parameter set, we can use a small-gain theorem for monotone systems from [7]. We reproduce the result as Theorem S2 in Section A.2 in

Supplementary Information. In particular, to verify stability using this small gain result, we consider the following system:

$$\begin{aligned}
\frac{d[\mathrm{HK}]}{dt} &= \beta_{\mathrm{HK}} - \delta[\mathrm{HK}] - k_{\mathrm{ap}}[\mathrm{HK}] + k_t(\mathrm{HK}_{\mathrm{tot}} - [\mathrm{HK}])[\mathrm{RR}] + k_{tc}(\mathrm{HK}_{\mathrm{tot}} - [\mathrm{HK}])[\mathrm{SR}],\\
\frac{d[\mathrm{RR}]}{dt} &= \beta_{\mathrm{RR}} - \delta[\mathrm{RR}] - k_t(\mathrm{HK}_{\mathrm{tot}} - [\mathrm{HK}])[\mathrm{RR}] + k_p[\mathrm{HK}](\mathrm{RR}_{\mathrm{tot}} - [\mathrm{RR}]),\\
\frac{d[\mathrm{SR}]}{dt} &= u - \delta[\mathrm{SR}] - k_{tc}(\mathrm{HK}_{\mathrm{tot}} - [\mathrm{HK}])[\mathrm{SR}] + k_{pc}[\mathrm{HK}]([\mathrm{SR}_{\mathrm{sum}}] - [\mathrm{SR}]),\\
\frac{d[\mathrm{SR}_{\mathrm{sum}}]}{dt} &= u - \delta[\mathrm{SR}_{\mathrm{sum}}],\\
y &= \mathrm{RR}_{\mathrm{tot}} - [\mathrm{RR}],\\
u &= Pk_{\mathrm{out}}([\mathrm{RR}]) = Pk_{\mathrm{out-max}}\frac{(y/K_{dr})^n}{(y/K_{dr})^n + 1},
\end{aligned} \tag{S19}$$

where we treat $\beta_{\mathrm{SR}}$ as an input $u$ to the signal sequestration motif (the open-loop system (4)) with an output $y = \mathrm{RR}_{\mathrm{tot}} - [\mathrm{RR}]$. Closing the loop is performed by the relation $\beta_{\mathrm{SR}} = Pk_{\mathrm{out}}([\mathrm{RR}])$. Note that the system (S19) is monotone with respect to the input $u$ and the feedback is negative if $y = \mathrm{RR}_{\mathrm{tot}} - [\mathrm{RR}]$ and the loop is closed with $u = Pk_{\mathrm{out-max}}\frac{(y/K_{dr})^n}{(y/K_{dr})^n+1}$. Proving stability is again performed using simulations.

#### B.4 Approximate Steady-State Computations

The closed-form steady-state computations can be reduced to solving four polynomial equations, which is not straightforward. Therefore, we resort to approximations and numerical computations. First, we derive an approximate closed-form steady-state expression for the sequestration mechanism model, which is valid under certain assumptions and was presented in the main text in Figure 2.

**Proposition S2** *1. Consider the phosphorylation sequestration motif (i.e., the system* (S16)*) under the following assumptions*

*(a) The total concentration of the response regulators is much larger than the concentration of the phosphorylated proteins:* $[\mathrm{RR_p}] \ll [\mathrm{RR}_{\mathrm{sum}}]$, $[\mathrm{SR_p}] \ll [\mathrm{SR}_{\mathrm{sum}}]$

*(b) The following holds:*

$$\delta + k_t\mathrm{RR}_{\mathrm{tot}} + k_{tc}\mathrm{SR}_{\mathrm{tot}} \gg \frac{k_{\mathrm{ap}}(k_t\mathrm{HK}_{\mathrm{tot}} + \delta)}{\delta + k_p\mathrm{HK}_{\mathrm{tot}}}.$$

*the steady state of* $[\mathrm{RR_p}]$ *can be approximated as follows:*

$$\boxed{[\mathrm{RR_p}]^* = \frac{k_{\mathrm{ap}}}{k_p}\cdot\frac{k_t\mathrm{RR}_{\mathrm{tot}}}{k_t\mathrm{RR}_{\mathrm{tot}} + k_{tc}\mathrm{SR}_{\mathrm{tot}} + \delta}\cdot\frac{k_p\mathrm{HK}_{\mathrm{tot}}}{k_p\mathrm{HK}_{\mathrm{tot}} + \delta}}. \tag{S20}$$

*Proof:* Since $[\mathrm{RR}] \approx [\mathrm{RR}_{\mathrm{sum}}] = \mathrm{RR}_{\mathrm{tot}}$, $[\mathrm{SR}] \approx [\mathrm{SR}_{\mathrm{sum}}] = \mathrm{SR}_{\mathrm{tot}}$, we have the following steady-state expressions for [HK] and $[\mathrm{HK_p}]$

$$\begin{aligned}
[\mathrm{HK_p}]^* &= \frac{k_{\mathrm{ap}}\mathrm{HK}_{\mathrm{tot}}}{\delta + k_{\mathrm{ap}} + k_t\mathrm{RR}_{\mathrm{tot}} + k_{tc}\mathrm{SR}_{\mathrm{tot}}},\\
[\mathrm{HK}]^* &= \frac{(\delta + k_{tc}\mathrm{SR}_{\mathrm{tot}} + k_t\mathrm{RR}_{\mathrm{tot}})\mathrm{HK}_{\mathrm{tot}}}{\delta + k_{\mathrm{ap}} + k_t\mathrm{RR}_{\mathrm{tot}} + k_{tc}\mathrm{SR}_{\mathrm{tot}}}.
\end{aligned}$$

Substituting this expression into the steady-state equation for $[\mathrm{RR_p}]$ we get:

$$\left(\delta + \frac{k_t k_{\mathrm{ap}}\mathrm{HK}_{\mathrm{tot}}}{\delta + k_{\mathrm{ap}} + k_t\mathrm{RR}_{\mathrm{tot}} + k_{tc}\mathrm{SR}_{\mathrm{tot}}} + \frac{k_p(\delta + k_t\mathrm{RR}_{\mathrm{tot}} + k_{tc}\mathrm{SR}_{\mathrm{tot}})\mathrm{HK}_{\mathrm{tot}}}{\delta + k_{\mathrm{ap}} + k_t\mathrm{RR}_{\mathrm{tot}} + k_{tc}\mathrm{SR}_{\mathrm{tot}}}\right)[\mathrm{RR_p}] = \frac{k_t k_{\mathrm{ap}}\mathrm{HK}_{\mathrm{tot}}}{\delta + k_{\mathrm{ap}} + k_t\mathrm{RR}_{\mathrm{tot}} + k_{tc}\mathrm{SR}_{\mathrm{tot}}}\mathrm{RR}_{\mathrm{tot}},$$

which can be simplified to

$$[\mathrm{RR_p}] = \frac{k_t \mathrm{RR_{tot}}}{(k_p \mathrm{HK_{tot}} + \delta)(\delta + k_t \mathrm{RR_{tot}} + k_{tc}\mathrm{SR_{tot}}) + k_t k_{\mathrm{ap}} \mathrm{HK_{tot}} + \delta k_{\mathrm{ap}}} k_{\mathrm{ap}} \mathrm{HK_{tot}} = \frac{k_t \mathrm{RR_{tot}}}{\delta + k_t \mathrm{RR_{tot}} + k_{tc}\mathrm{SR_{tot}} + \dfrac{k_{\mathrm{ap}}(k_t \mathrm{HK_{tot}} + \delta)}{k_p \mathrm{HK_{tot}} + \delta}} k_{\mathrm{ap}} \frac{\mathrm{HK_{tot}}}{k_p \mathrm{HK_{tot}} + \delta} \approx \frac{k_{\mathrm{ap}}}{k_p} \cdot \frac{k_t \mathrm{RR_{tot}}}{\delta + k_t \mathrm{RR_{tot}} + k_{tc}\mathrm{SR_{tot}}} \cdot \frac{k_p \mathrm{HK_{tot}}}{k_p \mathrm{HK_{tot}} + \delta}.$$

Finally, we have:

$$\boxed{[\mathrm{RR_p}] = \frac{k_{\mathrm{ap}}}{k_p} \cdot \frac{k_t \mathrm{RR_{tot}}}{k_t \mathrm{RR_{tot}} + k_{tc}\mathrm{SR_{tot}} + \delta} \cdot \frac{k_p \mathrm{HK_{tot}}}{k_p \mathrm{HK_{tot}} + \delta}}$$

■

These assumptions are fulfilled if, for example, $k_{\mathrm{ap}}$ is sufficiently small and $\mathrm{RR_{tot}}$ is much larger than $\mathrm{HK_{tot}}$. The first assumption is justified if the dissociation constant for the promoter initiating the output protein transcription is much lower than the total concentration of the response regulator protein. We are then interested in the regime when the concentration of the phosphorylated response regulators is much smaller than the total concentration of the response regulators.

In order to compute the steady-state of the closed loop system, we need to eliminate the variable $\mathrm{SR_{tot}}$ using the following relationship, which comes from the production of SR, where $P$ is the feedback strength:

$$\mathrm{SR_{tot}} = \frac{P k_{\mathrm{out-max}}/\delta([\mathrm{RR_p}]/K_{\mathrm{dr}})^n}{([\mathrm{RR_p}]/K_{\mathrm{dr}})^n + 1}. \tag{S21}$$

Hence, we need to solve the following equation:

$$[\mathrm{RR_p}] = \frac{a(([\mathrm{RR_p}]/K_{\mathrm{dr}})^n + 1)}{b([\mathrm{RR_p}]/K_{\mathrm{dr}})^n + 1}, \tag{S22}$$

where

$$a = \frac{k_{\mathrm{ap-max}}}{k_p} \frac{[\mathrm{I}]}{[\mathrm{I}] + K_{\mathrm{da}}} \frac{k_p \mathrm{HK_{tot}}}{k_p \mathrm{HK_{tot}} + \delta},$$
$$b = \frac{k_t \mathrm{RR_{tot}} + k_{tc} P k_{\mathrm{out-max}}/\delta}{k_t \mathrm{RR_{tot}}}.$$

The equation (S22) is hard to solve analytically unless $n = 1$, but can be solved numerically. Consider the function on the right hand side and letting $x = [\mathrm{RR_p}]/K_{\mathrm{dr}}$, we have

$$\frac{a(x^n + 1)}{bx^n + 1} = a/b + \frac{a - a/b}{bx^n + 1},$$

where $b \geq 1$ and hence $a - a/b \geq 0$. This this function is monotonically decreasing from $a$ for $[\mathrm{RR_p}] = 0$ to $a/b$ as $[\mathrm{RR_p}]$ goes to infinity. Therefore, there is a unique root to equation (S22), which implies that the system has a unique steady-state.

We can rewrite the model (S20) for the phosphorylation sequestration motif in a similar way to the RNAP and ribosome sharing model (13):

$$[\mathrm{RR_p}] = \frac{k_t}{\delta} \frac{k_{\mathrm{ap}}}{k_p} \frac{k_p \mathrm{HK_{tot}}}{k_p \mathrm{HK_{tot}} + \delta} \cdot \frac{\mathrm{RR_{tot}}}{\frac{k_t}{\delta}\mathrm{RR_{tot}} + \frac{k_{tc}}{\delta}\mathrm{SR_{tot}} + 1},$$

where $k_{\mathrm{ap}}$, $k_p$, $k_t$ are kinase autophosphorylation, response regulator phosphorylation and dephosphorylation rates, respectively; $\delta$ is the dilution rate; $\beta_{\mathrm{HK}}$, $\beta_{\mathrm{RR}}$, $\beta_{\mathrm{SR}}$, $\beta_{\mathrm{PH}}$ are HK, RR, SR, and PH production rates, respectively; and the other constants are computed as follows: $\mathrm{RR_{tot}} = \beta_{\mathrm{RR}}/\delta$, $\mathrm{SR_{tot}} = \beta_{\mathrm{SR}}/\delta$, $\mathrm{HK_{tot}} = \beta_{\mathrm{HK}}/\delta$, and $\mathrm{PH_{tot}} = \beta_{\mathrm{PH}}/\delta$ (see Sections B.2 and C.1). The constants $k_t/\delta$, $k_{tc}/\delta$ signify the resource usage by the two response regulators.

#### B.5 Frequency Domain Analysis

Since we perform the analysis around the steady-state, we can assume that $[\mathrm{HK_{sum}}] = \mathrm{HK_{tot}}$ and $[\mathrm{RR_{sum}}] = \mathrm{RR_{tot}}$. We separate the model into the process:

$$\begin{aligned}\frac{d[\mathrm{HK}]}{dt} &= \beta_{\mathrm{HK}} + (k_t(\mathrm{RR_{tot}} - [\mathrm{RR_p}]) + k_{tc}[\mathrm{SR}])(\mathrm{HK_{tot}} - [\mathrm{HK}]) - (k_{\mathrm{ap}} + \delta)[\mathrm{HK}],\\ \frac{d[\mathrm{RR_p}]}{dt} &= -\delta[\mathrm{RR_p}] + k_t(\mathrm{HK_{tot}} - [\mathrm{HK}])(\mathrm{RR_{tot}} - [\mathrm{RR_p}]) - k_p[\mathrm{HK}][\mathrm{RR_p}],\end{aligned} \tag{S23}$$

and the controller:

$$\begin{aligned}\frac{d[\mathrm{SR}]}{dt} &= h([\mathrm{RR_p}]) - \delta[\mathrm{SR}] - k_{tc}(\mathrm{HK_{tot}} - [\mathrm{HK}])[\mathrm{SR}] + k_{pc}[\mathrm{HK}]([\mathrm{SR_{sum}}] - [\mathrm{SR}]),\\ \frac{d[\mathrm{SR_{sum}}]}{dt} &= h([\mathrm{RR_p}]) - \delta[\mathrm{SR_{sum}}],\\ h(x) &= \frac{k_{\mathrm{out-max}}(x/K_{\mathrm{dr}})^n}{(x/K_{\mathrm{dr}})^n + 1}.\end{aligned} \tag{S24}$$

The process has one external input – the inducer concentration entering the equation through the autophosphorylation rate $k_{\mathrm{ap}}$, which we denote as $z$. The process also has a controlled input [SR] (denoted as $u$) and the outputs [HK] ($y_1$) and $[\mathrm{RR_p}]$ ($y_2$), while the controller has the output [SR] and the inputs [HK] and $[\mathrm{RR_p}]$.

Linearising the process around the steady-state $[\mathrm{RR_p}]^*$, $[\mathrm{HK}]^*$, $[\mathrm{SR}]^*$ gives:

$$\begin{aligned}\dot{x}_{s1} &= -(\delta + k_{\mathrm{ap}} + k_t(\mathrm{RR_{tot}} - [\mathrm{RR_p}]^*) + k_{tc}[\mathrm{SR}]^*)x_{s1} - k_t(\mathrm{HK_{tot}} - [\mathrm{HK}]^*)x_{s2}\\ &\qquad + k_{tc}(\mathrm{HK_{tot}} - [\mathrm{HK}]^*)u - \frac{\partial k_{\mathrm{ap}}}{\partial[\mathrm{I}]}[\mathrm{HK}]^* z\\ \dot{x}_{s2} &= -(k_t(\mathrm{RR_{tot}} - [\mathrm{RR_p}]^*) + k_p[\mathrm{RR_p}]^*)x_{s1} - (\delta + k_t(\mathrm{HK_{tot}} - [\mathrm{HK}]^*) + k_p[\mathrm{HK}]^*)x_{s2},\\ y_1 &= x_{s1},\\ y_2 &= x_{s2},\end{aligned}$$

where $x_{s1}$ is the deviation from the steady-state value $[\mathrm{HK}]^*$ and $x_{s2}$ is the deviation from the steady-state value $[\mathrm{RR_p}]^*$. We use the following short-hand notation for the process:

$$\begin{aligned}\frac{d}{dt}\begin{pmatrix}x_{s1}\\ x_{s2}\end{pmatrix} &= A_{P_{\mathrm{rr}}}\begin{pmatrix}x_{s1}\\ x_{s2}\end{pmatrix} + B_{P_{\mathrm{rr}}}\begin{pmatrix}u\\ z\end{pmatrix},\\ y &= C_{P_{\mathrm{rr}}}\begin{pmatrix}x_{s1}\\ x_{s2}\end{pmatrix},\end{aligned}$$

where

$$\begin{aligned}A_{P_{\mathrm{rr}}} &= \begin{pmatrix}-(\delta + k_{\mathrm{ap}} + k_t(\mathrm{RR_{tot}} - [\mathrm{RR_p}]^*) + k_{tc}[\mathrm{SR}]^*) & -k_t(\mathrm{HK_{tot}} - [\mathrm{HK}]^*)\\ -(k_t(\mathrm{RR_{tot}} - [\mathrm{RR_p}]^*) + k_p[\mathrm{RR_p}]^*) & -(\delta + k_t(\mathrm{HK_{tot}} - [\mathrm{HK}]^*) + k_p[\mathrm{HK}]^*)\end{pmatrix}\\ B_{P_{\mathrm{rr}}} &= \begin{pmatrix}k_{tc}(\mathrm{HK_{tot}} - [\mathrm{HK}]^*) & -\frac{\partial k_{\mathrm{ap}}}{\partial[\mathrm{I}]}[\mathrm{HK}]^*\\ 0 & 0\end{pmatrix} C_{P_{\mathrm{rr}}} = \begin{pmatrix}1 & 0\\ 0 & 1\end{pmatrix}.\end{aligned}$$

**Proposition S3** *The matrix $A_{P_{\mathrm{rr}}}$ is Hurwitz if*

$$\begin{aligned}&(\delta + k_{\mathrm{ap}} + k_t(\mathrm{RR_{tot}} - [\mathrm{RR_p}]^*) + k_{tc}[\mathrm{SR}]^*)(\delta + k_t(\mathrm{HK_{tot}} - [\mathrm{HK}]^*) + k_p[\mathrm{HK}]^*) >\\ &\qquad k_t(\mathrm{HK_{tot}} - [\mathrm{HK}]^*)(k_t(\mathrm{RR_{tot}} - [\mathrm{RR_p}]^*) + k_p[\mathrm{RR_p}]^*).\end{aligned} \tag{S25}$$

*Proof:* The matrix $A_{P_{\text{rr}}}$ is two by two therefore it is stable if its trace is negative and determinant is positive. Since the elements on the diagonal are negative, we only have to check the sign of the determinant, which leads to the condition (S25). ■

Note that the condition (S25) can be made parameter independent while sacrificing some generality. Indeed, we have

$$(\delta + k_{\text{ap}} + k_t(\text{RR}_{\text{tot}} - [\text{RR}_\text{p}]^*) + k_{tc}[\text{SR}]^*)(\delta + k_t(\text{HK}_{\text{tot}} - [\text{HK}]^*) + k_p[\text{HK}]^*) > \\ k_t(\text{RR}_{\text{tot}} - [\text{RR}_\text{p}]^*)(k_t(\text{HK}_{\text{tot}} - [\text{HK}]^*) + k_p[\text{HK}]^*).$$

Therefore, if

$$k_t(\text{RR}_{\text{tot}} - [\text{RR}_\text{p}]^*)(k_t(\text{HK}_{\text{tot}} - [\text{HK}]^*) + k_p[\text{HK}]^*) > k_t(\text{HK}_{\text{tot}} - [\text{HK}]^*)(k_t(\text{RR}_{\text{tot}} - [\text{RR}_\text{p}]^*) + k_p[\text{RR}_\text{p}]^*),$$

we also have stability. We have the following chain of equivalent identities:

$$k_pk_t(\text{RR}_{\text{tot}} - [\text{RR}_\text{p}]^*)[\text{HK}]^* > k_pk_t(\text{HK}_{\text{tot}} - [\text{HK}]^*)[\text{RR}_\text{p}]^*,$$
$$\Updownarrow$$
$$\text{RR}_{\text{tot}}[\text{HK}]^* - [\text{RR}_\text{p}]^*[\text{HK}]^* > \text{HK}_{\text{tot}}[\text{RR}_\text{p}]^* - [\text{HK}]^*[\text{RR}_\text{p}]^*$$
$$\Updownarrow$$
$$\frac{[\text{HK}]^*}{\text{HK}_{\text{tot}}} > \frac{[\text{RR}_\text{p}]^*}{\text{RR}_{\text{tot}}}$$
$$\Updownarrow$$
$$1 > \frac{[\text{HK}_\text{p}]^*}{\text{HK}_{\text{tot}}} + \frac{[\text{RR}_\text{p}]^*}{\text{RR}_{\text{tot}}}.$$

Therefore, if the proportion of not phosphorylated HK in the steady state is lower than the proportion of the phosphorylated RR then the open-loop system is globally stable. Equivalently, if the sum of proportions of phosphorylated HK and RR are less than one, the system is stable.

The controller dynamics are as follows:

$$\frac{d[\text{SR}]}{dt} = h([\text{RR}_\text{p}]) - \delta[\text{SR}] - k_{tc}[\text{HK}_\text{p}][\text{SR}] + k_{pc}[\text{HK}]([\text{SR}_{\text{sum}}] - [\text{SR}]),$$
$$\frac{d[\text{SR}_{\text{sum}}]}{dt} = h([\text{RR}_\text{p}]) - \delta[\text{SR}_{\text{sum}}],$$
$$h(x) = \frac{Pk_{\text{out}-\text{max}}(x/K_{dr})^n}{(x/K_{dr})^n + 1}.$$

The linearised dynamics of the controller around the steady-state $[\text{RR}_\text{p}]^*$, $[\text{HK}]^*$, $[\text{SR}]^*$, $[\text{SR}_{\text{sum}}]^*$ take the following form:

$$\dot{x}_{k1} = -(\delta + k_{tc}(\text{HK}_{\text{tot}} - [\text{HK}]^*) + k_{pc}[\text{HK}]^*)x_{k1} + k_{pc}[\text{HK}]^*x_{k2} \\ + (k_{pc}([\text{SR}_{\text{sum}}]^* - [\text{SR}]^*) + k_{tc}[\text{SR}]^*)y_1 + \frac{\partial h([\text{RR}_\text{p}]^*)}{\partial [\text{RR}_\text{p}]^*}y_2, \tag{S26}$$

$$\dot{x}_{k2} = -\delta x_{k2} + \frac{\partial h([\text{RR}_\text{p}]^*)}{\partial [\text{RR}_\text{p}]^*}y_2, \tag{S27}$$

$$u = x_{k1}, \tag{S28}$$

where $x_{k1}$ is the deviation from the steady-state value $[\text{SR}]^*$ and $x_{k2}$ is the deviation from the steady-state value $[\text{SR}_{\text{sum}}]^*$. The transfer function can be computed as follows:

$$K_{\text{uy}} = \begin{pmatrix} K_{\text{uy}_1} & K_{\text{uy}_2} \end{pmatrix} = \begin{pmatrix} \dfrac{b_1}{s+a_1} & \dfrac{b_2}{s+a_3}\dfrac{s+a_2+a_3}{s+a_1} \end{pmatrix},$$

where

$$
\begin{aligned}
a_1 &= \delta + k_{tc}(\mathrm{HK_{tot}} - [\mathrm{HK}]^*) + k_{pc}[\mathrm{HK}]^*,\\
a_2 &= k_{pc}[\mathrm{HK}]^*,\\
a_3 &= \delta,\\
b_1 &= k_{pc}([\mathrm{SR_{sum}}]^* - [\mathrm{SR}]^*) + k_{tc}[\mathrm{SR}]^*,\\
b_2 &= \frac{\partial h([\mathrm{RR_p}]^*)}{\partial [\mathrm{RR_p}]^*}.
\end{aligned}
$$

Note that $a_2 + a_3 \approx a_1$, if the concentration of phosphorylated [HK] in the steady state is low. Therefore, there is a near pole-zero cancellation, which in this case is not detrimental for the behaviour of the closed loop, since the poles are stable and the slowest dynamics are defined by the pole at $a_3 = \delta$. This means that the controller is essentially a low-pass filter with the major limitation in the form of the dilution/degradation rate of the system.

In order to compute the sensitivity function, as discussed in Section A.3, we perturb the measured output ($y_2$) corresponding to [$\mathrm{RR_p}$] by the noise term $w$, and we only need to consider the transfer function of the process from $u$ to $y_1$ and $y_2$. As we do not perturb the input $y_1$, we compute the sensitivity function as discussed in Section A.3, but take only its $(2, 2)$-entry.

The state-space of the process (and the controller) has a specific sign pattern, meaning the sign of the entry does not change for all possible steady-state values. In particular we have the following sign-pattern:

$$
\left[\begin{array}{c|c} A_P & B_P \\ \hline C_P & 0 \end{array}\right] \rightarrow \left[\begin{array}{cc|c} - & - & + \\ - & - & 0 \\ \hline + & + & 0 \end{array}\right].
$$

This sign pattern implies that the controller $K_{\mathrm{uy_1}}$ acts as a positive feedback on the system $G_{\mathrm{y_1u}}$, while the controller $K_{\mathrm{uy_2}}$ forms a negative feedback loop with $G_{\mathrm{y_2u}}$. We can view, however, the controller $K_{\mathrm{uy_1}}$ and the process $G_{\mathrm{y_1u}}$ as retroactivity appearing while connecting a negative feedback controller to the loop. In our case, the rectroactive components result in minimum-phase transfer functions, therefore, they do not have a detrimental effect on stability or performance.

### C Dephosphorylation Enhancement Motif Modelling

#### C.1 Basic Model

We took the chemical reactions for the wild-type model (S10) and assumed the following chemical reactions:

$$
\begin{aligned}
&\mathrm{RR_p} + \mathrm{PH} \xrightarrow{k_{pc}} \mathrm{PH} + \mathrm{RR},\\
&\mathrm{PH} \xrightarrow{\delta} \emptyset, \qquad\qquad \emptyset \xrightarrow{\beta_{\mathrm{PH}}} \mathrm{PH}.
\end{aligned} \tag{S29}
$$

Assuming $\beta_{\mathrm{PH}} = Pk_{\mathrm{out}}$ results in the following set of differential equations:

$$
\begin{aligned}
\frac{d[\mathrm{HK}]}{dt} &= \beta_{\mathrm{HK}} - \delta[\mathrm{HK}] - k_{\mathrm{ap}}[\mathrm{HK}] + k_t[\mathrm{HK_p}][\mathrm{RR}],\\
\frac{d[\mathrm{HK_p}]}{dt} &= -\delta[\mathrm{HK_p}] + k_{\mathrm{ap}}[\mathrm{HK}] - k_t[\mathrm{HK_p}][\mathrm{RR}],\\
\frac{d[\mathrm{RR}]}{dt} &= \beta_{\mathrm{RR}} - \delta[\mathrm{RR}] - k_t[\mathrm{HK_p}][\mathrm{RR}] + k_p[\mathrm{HK}][\mathrm{RR_p}] + k_{pc}[\mathrm{PH}][\mathrm{RR_p}],\\
\frac{d[\mathrm{RR_p}]}{dt} &= -\delta[\mathrm{RR_p}] + k_t[\mathrm{HK_p}][\mathrm{RR}] - k_p[\mathrm{HK}][\mathrm{RR_p}] - k_{pc}[\mathrm{PH}][\mathrm{RR_p}],\\
\frac{d[\mathrm{PH}]}{dt} &= Pk_{\mathrm{out}}([\mathrm{RR_p}]) - \delta[\mathrm{PH}],\\
\frac{d[\mathrm{Output}]}{dt} &= k_{\mathrm{out}}([\mathrm{RR_p}]) - \delta[\mathrm{Output}].
\end{aligned} \tag{S30}
$$

#### C.2 Analysis of the Dephosphorylation Enhancement Model

We consider first the model of the sequestration mechanism:

$$\begin{aligned}
\frac{d[\mathrm{HK_p}]}{dt} &= -\delta[\mathrm{HK_p}] + k_{\mathrm{ap}}([\mathrm{HK_{sum}}] - [\mathrm{HK_p}]) - k_t[\mathrm{HK_p}]([\mathrm{RR_{sum}}] - [\mathrm{RR_p}]),\\
\frac{d[\mathrm{RR_p}]}{dt} &= -\delta[\mathrm{RR_p}] + k_t[\mathrm{HK_p}]([\mathrm{RR_{sum}}] - [\mathrm{RR_p}]) - (k_p[\mathrm{HK}] + k_{pc}[\mathrm{PH}])[\mathrm{RR_p}],\\
\frac{d[\mathrm{PH}]}{dt} &= \beta_{\mathrm{PH}} - \delta[\mathrm{PH}],\\
\frac{d[\mathrm{HK_{sum}}]}{dt} &= \beta_{\mathrm{HK}} - \delta[\mathrm{HK_{sum}}],\\
\frac{d[\mathrm{RR_{sum}}]}{dt} &= \beta_{\mathrm{RR}} - \delta[\mathrm{RR_{sum}}],
\end{aligned}$$

which is forward-invariant with respect to the set

$$\begin{aligned}
\mathcal{S}_T = \Big\{ \big( [\mathrm{HK}] \quad [\mathrm{RR}] \quad [\mathrm{PH}] \quad [\mathrm{HK_{sum}}] \quad [\mathrm{RR_{sum}}] \big)^T \Big| \\
0 \le [\mathrm{HK}] \le \mathrm{HK_{tot}}, 0 \le [\mathrm{PH}] \le \mathrm{PH_{tot}}, \\
0 \le [\mathrm{RR}] \le \mathrm{RR_{tot}}, 0 \le [\mathrm{HK_{sum}}] \le \mathrm{HK_{tot}}, 0 \le [\mathrm{RR_{sum}}] \le \mathrm{RR_{tot}} \Big\}.
\end{aligned}$$

As above, if we assume that $[\mathrm{HK_{sum}}]$ equals to $\mathrm{HK_{tot}}$ at time zero, then we can produce a model, monotone with respect to all parameters, but $k_t$ and $\mathrm{HK_{tot}}$. Stability verification of this model can be performed in a similar manner to the phosphorylation sequestration model.

#### C.3 Approximate Steady-State

Here, we derive the steady-state model of the dephosphorylation enhancement mechanism presented in the main text in Figure 2. Recall that we consider the following system in the main text:

$$\begin{aligned}
\frac{d[\mathrm{HK}]}{dt} &= \beta_{\mathrm{HK}} - \delta[\mathrm{HK}] - k_{\mathrm{ap}}[\mathrm{HK}] + k_t[\mathrm{HK_p}][\mathrm{RR}],\\
\frac{d[\mathrm{HK_p}]}{dt} &= -k_t[\mathrm{HK_p}][\mathrm{RR}] + k_{\mathrm{ap}}[\mathrm{HK}] - \delta[\mathrm{HK_p}],\\
\frac{d[\mathrm{RR}]}{dt} &= \beta_{\mathrm{RR}} - \delta[\mathrm{RR}] - k_t[\mathrm{HK_p}][\mathrm{RR}] + k_p[\mathrm{HK}][\mathrm{RR_p}] + k_{pc}[\mathrm{PH}][\mathrm{RR_p}],\\
\frac{d[\mathrm{RR_p}]}{dt} &= -\delta[\mathrm{RR_p}] + k_t[\mathrm{HK_p}][\mathrm{RR}] - k_p[\mathrm{HK}][\mathrm{RR_p}] - k_{pc}[\mathrm{PH}][\mathrm{RR_p}],\\
\frac{d[\mathrm{PH}]}{dt} &= \beta_{\mathrm{PH}} - \delta[\mathrm{PH}],\\
\frac{d[\mathrm{Output}]}{dt} &= k_{\mathrm{out}}([\mathrm{RR_p}]) - \delta[\mathrm{Output}],
\end{aligned} \tag{S31}$$

**Proposition S4** *2. Consider the dephosphorylation enhacement motif (i.e., the system* (S31)*). If the following holds:*

$$k_t\mathrm{RR_{tot}} + \delta \gg k_{\mathrm{ap}} \frac{\delta + k_{pc}\mathrm{PH_{tot}} + k_t\mathrm{HK_{tot}}}{\delta + k_{pc}\mathrm{PH_{tot}} + k_p\mathrm{HK_{tot}}},$$

*then the steady state of* $[\mathrm{RR_p}]$ *can be approximated as follows:*

$$\boxed{[\mathrm{RR_p}]^* = \frac{k_{\mathrm{ap}}}{k_p} \cdot \frac{k_t\mathrm{RR_{tot}}}{k_t\mathrm{RR_{tot}} + \delta} \cdot \frac{k_p\mathrm{HK_{tot}}}{\delta + k_{pc}\mathrm{PH_{tot}} + k_p\mathrm{HK_{tot}}}} \tag{S32}$$

*Proof:* By setting the derivatives to zero we obtain the following relations:

$$
\begin{aligned}
[\mathrm{PH}] &= \mathrm{PH}_{\mathrm{tot}}, \\
[\mathrm{RR}_{\mathrm{sum}}] &= \mathrm{RR}_{\mathrm{tot}}, \\
[\mathrm{HK}_{\mathrm{p}}] &= \frac{k_{\mathrm{ap}}\mathrm{HK}_{\mathrm{tot}}}{\delta + k_{\mathrm{ap}} + k_t(\mathrm{RR}_{\mathrm{tot}} - \mathrm{RR}_{\mathrm{p}})} \\
[\mathrm{HK}] &= \frac{(k_t(\mathrm{RR}_{\mathrm{tot}} - \mathrm{RR}_{\mathrm{p}}) + \delta)\mathrm{HK}_{\mathrm{tot}}}{\delta + k_{\mathrm{ap}} + k_t(\mathrm{RR}_{\mathrm{tot}} - \mathrm{RR}_{\mathrm{p}})} \\
[\mathrm{RR}_{\mathrm{p}}] &= \frac{k_t[\mathrm{HK}_{\mathrm{p}}]\mathrm{RR}_{\mathrm{tot}}}{\delta + k_p\mathrm{HK}_{\mathrm{tot}} + k_{pc}[\mathrm{PH}] + (k_t - k_p)[\mathrm{HK}_{\mathrm{p}}]}
\end{aligned}
$$

Combining the relations yields the following chain of equalities:

$$
\begin{aligned}
&[\mathrm{RR}_{\mathrm{p}}]\left((\delta + k_{pc}\mathrm{PH}_{\mathrm{tot}} + k_p\mathrm{HK}_{\mathrm{tot}})(k_t(\mathrm{RR}_{\mathrm{tot}} - [\mathrm{RR}_{\mathrm{p}}]) + \delta + k_{\mathrm{ap}})\right. \\
&\qquad \left. + (k_t - k_p)k_{\mathrm{ap}}\mathrm{HK}_{\mathrm{tot}}\right) - k_t k_{\mathrm{ap}}\mathrm{RR}_{\mathrm{tot}}\mathrm{HK}_{\mathrm{tot}} = \\
&- [\mathrm{RR}_{\mathrm{p}}]^2(\delta + k_{pc}\mathrm{PH}_{\mathrm{tot}} + k_p\mathrm{HK}_{\mathrm{tot}})k_t + [\mathrm{RR}_{\mathrm{p}}]((\delta + k_{pc}\mathrm{PH}_{\mathrm{tot}} + k_p\mathrm{HK}_{\mathrm{tot}})(k_t\mathrm{RR}_{\mathrm{tot}} + \delta + k_{\mathrm{ap}}) \\
&\qquad + (k_t - k_p)k_{\mathrm{ap}}\mathrm{HK}_{\mathrm{tot}}) - k_t k_{\mathrm{ap}}\mathrm{RR}_{\mathrm{tot}}\mathrm{HK}_{\mathrm{tot}} = \\
&- [\mathrm{RR}_{\mathrm{p}}]^2 k_t + [\mathrm{RR}_{\mathrm{p}}]\left(\mathrm{RR}_{\mathrm{tot}}k_t + \delta + k_{\mathrm{ap}} + (k_t - k_p)\frac{k_{\mathrm{ap}}\mathrm{HK}_{\mathrm{tot}}}{\delta + k_{pc}\mathrm{PH}_{\mathrm{tot}} + k_p\mathrm{HK}_{\mathrm{tot}}}\right) \\
&\qquad - k_t\frac{k_{\mathrm{ap}}\mathrm{RR}_{\mathrm{tot}}\mathrm{HK}_{\mathrm{tot}}}{\delta + k_{pc}\mathrm{PH}_{\mathrm{tot}} + k_p\mathrm{HK}_{\mathrm{tot}}} = 0,
\end{aligned}
$$

which has one physically meaningful root:

$$
[\mathrm{RR}_{\mathrm{p}}] = \frac{(k_t\mathrm{RR}_{\mathrm{tot}} + \delta)}{2k_t}\left(z - \sqrt{z^2 - \frac{4k_t^2\mathrm{RR}_{\mathrm{tot}}k_{\mathrm{ap}}\mathrm{HK}_{\mathrm{tot}}}{(\delta + k_{\mathrm{pc}}\mathrm{PH}_{\mathrm{tot}} + k_{\mathrm{p}}\mathrm{HK}_{\mathrm{tot}})(k_{\mathrm{t}}\mathrm{RR}_{\mathrm{tot}} + \delta)^2}}\right),
$$

where

$$
z = \frac{1}{k_t\mathrm{RR}_{\mathrm{tot}} + \delta}\left(\mathrm{RR}_{\mathrm{tot}}k_t + \delta + k_{\mathrm{ap}} + (k_t - k_p)\frac{k_{\mathrm{ap}}\mathrm{HK}_{\mathrm{tot}}}{\delta + k_{pc}\mathrm{PH}_{\mathrm{tot}} + k_p\mathrm{HK}_{\mathrm{tot}}}\right).
$$

Indeed, it can be verified that the root

$$
[\mathrm{RR}_{\mathrm{p}}] = \frac{(k_t\mathrm{RR}_{\mathrm{tot}} + \delta)}{2k_t}\left(z + \sqrt{z^2 - \frac{4k_t^2\mathrm{RR}_{\mathrm{tot}}k_{\mathrm{ap}}\mathrm{HK}_{\mathrm{tot}}}{(\delta + k_{\mathrm{pc}}\mathrm{PH}_{\mathrm{tot}} + k_{\mathrm{p}}\mathrm{HK}_{\mathrm{tot}})(k_t\mathrm{RR}_{\mathrm{tot}} + \delta)^2}}\right)
$$

leads to $[\mathrm{RR}_{\mathrm{p}}] \geq \mathrm{RR}_{\mathrm{tot}}$, which violates our constraints. Note that $z \approx 1$ if

$$
k_t\mathrm{RR}_{\mathrm{tot}} + \delta \gg \left| k_{\mathrm{ap}} + (k_t - k_p)\frac{k_{\mathrm{ap}}\mathrm{HK}_{\mathrm{tot}}}{\delta + k_{pc}\mathrm{PH}_{\mathrm{tot}} + k_p\mathrm{HK}_{\mathrm{tot}}} \right| = k_{\mathrm{ap}}\frac{\delta + k_{pc}\mathrm{PH}_{\mathrm{tot}} + k_p\mathrm{HK}_{\mathrm{tot}}}{\delta + k_{pc}\mathrm{PH}_{\mathrm{tot}} + k_p\mathrm{HK}_{\mathrm{tot}}},
$$

which also leads to

$$
1 \gg \frac{k_{\mathrm{ap}}k_t\mathrm{HK}_{\mathrm{tot}}}{(\delta + k_{pc}\mathrm{PH}_{\mathrm{tot}} + k_p\mathrm{HK}_{\mathrm{tot}})(k_t\mathrm{RR}_{\mathrm{tot}} + \delta)}.
$$

Therefore, we also have

$$
1 \gg \frac{4k_t\mathrm{RR}_{\mathrm{tot}}}{(k_{\mathrm{t}}\mathrm{RR}_{\mathrm{tot}} + \delta)}\frac{k_{\mathrm{ap}}k_t\mathrm{HK}_{\mathrm{tot}}}{(\delta + k_{\mathrm{pc}}\mathrm{PH}_{\mathrm{tot}} + k_{\mathrm{p}}\mathrm{HK}_{\mathrm{tot}})(k_{\mathrm{t}}\mathrm{RR}_{\mathrm{tot}} + \delta)}.
$$

Now using that $z \approx 1$ and the Taylor series expansion

$$
\sqrt{1 - x} \approx 1 - \frac{1}{2}x,
$$

we obtain the following expression:

$$[\mathrm{RR_p}] = \frac{(k_t\mathrm{RR_{tot}}+\delta)}{2k_t}\cdot\frac{2k_t^2\mathrm{RR_{tot}}k_{\mathrm{ap}}\mathrm{HK_{tot}}}{(\delta+k_{\mathrm{pc}}\mathrm{PH_{tot}}+k_{\mathrm{p}}\mathrm{HK_{tot}})(k_t\mathrm{RR_{tot}}+\delta)^2} = \frac{k_{\mathrm{ap}}}{k_p}\cdot\frac{k_t\mathrm{RR_{tot}}}{k_{\mathrm{t}}\mathrm{RR_{tot}}+\delta}\cdot\frac{k_p\mathrm{HK_{tot}}}{\delta+k_{\mathrm{pc}}\mathrm{PH_{tot}}+k_{\mathrm{p}}\mathrm{HK_{tot}}}.$$

■

To compute the steady-state of the closed system we need to solve (S32) and

$$\mathrm{PH_{tot}} = \frac{Pk_{\mathrm{out-max}}/\delta([\mathrm{RR_p}]/K_{\mathrm{dr}})^n}{([\mathrm{RR_p}]/K_{\mathrm{dr}})^n+1} \tag{S33}$$

for $\mathrm{PH_{tot}}$ and $[\mathrm{RR_p}]$. Note that the solution is necessarily unique.

#### C.4 Frequency Domain Analysis

For frequency domain analysis we separate the system into the process and the controller in a similar way to the phosphorylation sequestration motif: We consider the process

$$\begin{aligned}
\frac{d[\mathrm{HK}]}{dt} &= \beta_{\mathrm{HK}} - (k_{\mathrm{ap}}+\delta)[\mathrm{HK}] + k_t(\mathrm{HK_{tot}}-[\mathrm{HK}])(\mathrm{RR_{tot}}-[\mathrm{RR_p}]),\\
\frac{d[\mathrm{RR_p}]}{dt} &= -\delta[\mathrm{RR_p}] + k_t(\mathrm{HK_{tot}}-[\mathrm{HK}])(\mathrm{RR_{tot}}-[\mathrm{RR_p}]) - (k_p[\mathrm{HK}]+k_{pc}[\mathrm{PH}])[\mathrm{RR_p}],
\end{aligned}$$

where we treat [PH] as the input and the phosphorylated RR as the output. The controller equations are

$$\frac{d[\mathrm{PH}]}{dt} = h([\mathrm{RR_p}]) - \delta[\mathrm{PH}], \tag{S34}$$

$$h(x) = \frac{k_{\mathrm{out-max}}(x/K_{dr})^n}{(x/K_{dr})^n+1}, \tag{S35}$$

where we treat $\mathrm{RR_p}$ as an input. We linearise the system around the steady-state $[\mathrm{RR_p}]^*$, $[\mathrm{HK}]^*$, $[\mathrm{PH}]^*$:

$$\begin{aligned}
\dot{x}_{s1} &= -(\delta+k_{\mathrm{ap}}+k_t(\mathrm{RR_{tot}}-[\mathrm{RR_p}]^*))x_{s1} - k_t(\mathrm{HK_{tot}}-[\mathrm{HK}]^*)x_{s2},\\
\dot{x}_{s2} &= -(k_t(\mathrm{RR_{tot}}-[\mathrm{RR_p}]^*)+k_p[\mathrm{RR_p}]^*)x_{s1},\\
&\qquad -(\delta+k_t(\mathrm{HK_{tot}}-[\mathrm{HK}]^*)+k_p[\mathrm{HK}]^*+k_{pc}[\mathrm{PH}]^*)x_{s2} - k_{pc}[\mathrm{RR_p}]^*u,\\
y &= x_{s2},
\end{aligned}$$

where $x_{s1}$ and $x_{s2}$ are the deviations from the steady-states $[\mathrm{HK}]^*$ and $[\mathrm{RR_p}]^*$, respectively. We use the following short-hand notation for the process:

$$\begin{aligned}
\frac{d}{dt}\begin{pmatrix}x_{s1}\\x_{s2}\end{pmatrix} &= A_{\mathrm{ph}}\begin{pmatrix}x_{s1}\\x_{s2}\end{pmatrix} + B_{\mathrm{ph}}u,\\
y &= C_{\mathrm{ph}}\left(x_{s2}\right),
\end{aligned}$$

where

$$A_{\mathrm{ph}} = \begin{pmatrix}-(\delta+k_{\mathrm{ap}}+k_t(\mathrm{RR_{tot}}-[\mathrm{RR_p}]^*)) & -k_t(\mathrm{HK_{tot}}-[\mathrm{HK}]^*)\\ -(k_t(\mathrm{RR_{tot}}-[\mathrm{RR_p}]^*)+k_p[\mathrm{RR_p}]^*) & -(\delta+k_t(\mathrm{HK_{tot}}-[\mathrm{HK}]^*)+k_p[\mathrm{HK}]^*+k_{pc}[\mathrm{PH}]^*)\end{pmatrix},$$

$$B_{\mathrm{ph}} = \begin{pmatrix}0\\ -k_{pc}[\mathrm{RR_p}]^*\end{pmatrix}, C_{\mathrm{ph}} = \begin{pmatrix}0 & 1\end{pmatrix}.$$

We obtain the following transfer function

$$G_{\rm ph}(s) = \frac{b(s - a_{11})}{(s - a_{11})(s - a_{22}) - a_{12}a_{21}},$$

where $a_{ij}$ are the entries of the matrix $A_{\rm ph}$ and $b = -k_{pc}[{\rm RR_p}]^*$.

The controller has linear dynamics and only the input is nonlinear. We only reproduce the transfer function here, which is as follows:

$$K_{\rm ph}(s) = \frac{\partial h([{\rm RR_p}]^*)}{\partial [{\rm RR_p}]^*} \cdot \frac{1}{s + \delta}.$$

This controller is the simplest low pass filter with a pole at the dilution rate.

### D Additional Numerical Simulations for the Dichotomous Feedback Mechanisms

#### D.1 Parameter Values

In order to analyse our theoretical model we pick the parameter values for the EnvZ (or Taz) - OmpR two-component system, which is a model two-component system that is well-studied in the literature. The parameter values vary depending on the study considered. The copy numbers of RR and HK have been reported to be in the range $100 - 2000$ and $3000 - 6000$, respectively [12–14], which corresponds to $0.167 - 3.3$ and $4.98 - 9.96$ [$\mu$M] in *E. Coli.* The autophosphorylation rate of Taz has been reported to be in the range $1.2 \cdot 10^{-1}$–6 [1/min] [14,15], the OmpR phosphorylation and dephosphorylation rates were measured between $6 - 6.21 \cdot 10^3$ [$1/(\mu$M min)] [14,16], and between $1.7 \cdot 10^{-1}$–3 [$1/(\mu$M min)] [12,14], respectively. The dissociation constant for *pOmpC* promoter was reported as $3.14 \cdot 10^{-2}$ [$\mu$M] [17]. The dilution rate $\delta$ was calculated by assuming cell division time of 30 [min] [18]. The *pOmpC* promoter has three binding cites, to which the dimers of the phosphorylated OmpR can bind, therefore the Hill coefficient for the system should be set to 6 [19]. However, in order to keep the results less specific to OmpR we tested the Hill coefficient equal to four. The parameters for autophosphorylation of HK, the expression of the output protein were taken from a reasonable parameter range. In the main text we took the parameter values from Table S1.

Table S1: Parameter values for theoretical analysis

| parameter | value/interval | units |
|---|---|---|
| $\delta$ | 0.0234 | 1/min |
| $\mathrm{HK_{tot}}$ | 0.167 | $\mu$M |
| $\mathrm{RR_{tot}}$ | 6 | $\mu$M |
| $\beta_{\rm HK}$ | $\delta\mathrm{HK_{tot}}$ | 1/min |
| $\beta_{\rm RR}$ | $\delta\mathrm{RR_{tot}}$ | 1/min |
| $k_t$ | $6.12 \cdot 10^3$ | 1/(min $\mu$M) |
| $k_p$ | $4 \cdot 10^{-3}$ | 1/(min $\mu$M) |
| $k_{tc}$ | $k_t$ | 1/(min $\mu$M) |
| $k_{pc}$ | $k_p$ | 1/(min $\mu$M) |
| $K_{\rm dr}$ | $3.14 \cdot 10^{-2}$ | $\mu$M |
| $K_{\rm da}$ | 2 | mM |
| $k_{\rm out-max}$ | 1 | 1/min |
| $k_{\rm ap-max}$ | 0.02 | 1/min |
| $P$ | [0, 1] | dimensionless |
| $n$ | 4 | dimensionless |

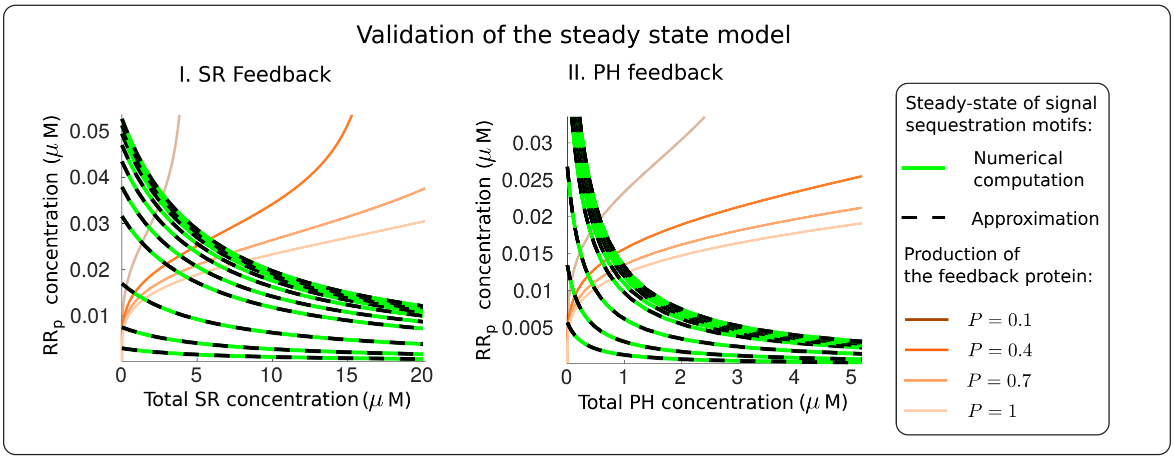

Figure S2: Numerical computations of the steady-states of the phosphorylation sequestration and the dephosphorylation enhancement mechanisms (black dashed curves), their analytical approximations in (S36, S38) (green curves) for different concentration of inducer in $[0, 10]$ [mM], and the curves of the feedback proteins production computes as in (S37, S39). The intersection points of the curves (S37), and (S36) (respectively, the curves (S39) and (S38)) are the steady-states of the SR feedback system (respectively, the PH feedback system). The numerical computations confirm that our steady-state models are accurate for the considered parameter range.

#### D.2 The Verification of the Steady-State Models

In order to distinguish between the two dichotomous feedback mechanisms, we refer to the feedback based on phosphorylation sequestration as SR feedback and to the feedback based on dephosphorylation enhancement as PH feedback. We verified our steady-state models using the parameter values from Table D.1 and the inducer concentration in the range $[0, 10]$ [mM], but we have used $k_p = 1.764 \cdot 10^{-1}$ $[1/(\mu\text{M}\,\text{min})]$. We computed the values of $[\text{RR}_\text{p}]$ for different inducer concentrations (note that $k_\text{ap}$ depends on the inducer concentration) and total concentrations of the protein SR ($\text{SR}_\text{tot}$) using the expression:

$$[\text{RR}_\text{p}] = \frac{k_\text{ap}}{k_p} \cdot \frac{k_t\text{RR}_\text{tot}}{k_t\text{RR}_\text{tot} + \delta} \cdot \frac{k_p\text{HK}_\text{tot}}{k_p\text{HK}_\text{tot} + \delta} \cdot \frac{1}{1 + \frac{k_{tc}}{k_t\text{RR}_\text{tot}+\delta}\text{SR}_\text{tot}}, \tag{S36}$$

and the values of $\text{SR}_\text{tot}$ depending on the concentration of $[\text{RR}_\text{p}]$ as follows:

$$\text{SR}_\text{tot} = \frac{Pk_{\text{out}-\max}/\delta([\text{RR}_\text{p}]/K_\text{dr})^6}{([\text{RR}_\text{p}]/K_\text{dr})^6 + 1}. \tag{S37}$$

As we discussed above the intersections of these curves are approximations of the steady states of the closed loop system. For the phosphatase feedback system, we performed similar computations using formulas:

$$[\text{RR}_\text{p}] = \frac{k_\text{ap}}{k_p} \cdot \frac{k_t\text{RR}_\text{tot}}{k_t\text{RR}_\text{tot} + \delta} \cdot \frac{k_p\text{HK}_\text{tot}}{k_p\text{HK}_\text{tot} + \delta} \cdot \frac{1}{1 + \frac{k_{pc}}{k_p\text{HK}_\text{tot}+\delta}\text{PH}_\text{tot}}, \tag{S38}$$

$$\text{PH}_\text{tot} = \frac{Pk_{\text{out}-\max}/\delta([\text{RR}_\text{p}]/K_\text{dr})^6}{([\text{RR}_\text{p}]/K_\text{dr})^6 + 1}. \tag{S39}$$

The computational results depicted in Figures S2.I and S2.II confirm the validity of the derived state-space models and the claimed similarity between the feedback mechanisms. While these figures were obtained with $k_{tc} = k_t$ and $k_{pc} = k_p$, we observed similar results when we varied $k_{tc}$ and $k_{pc}$.

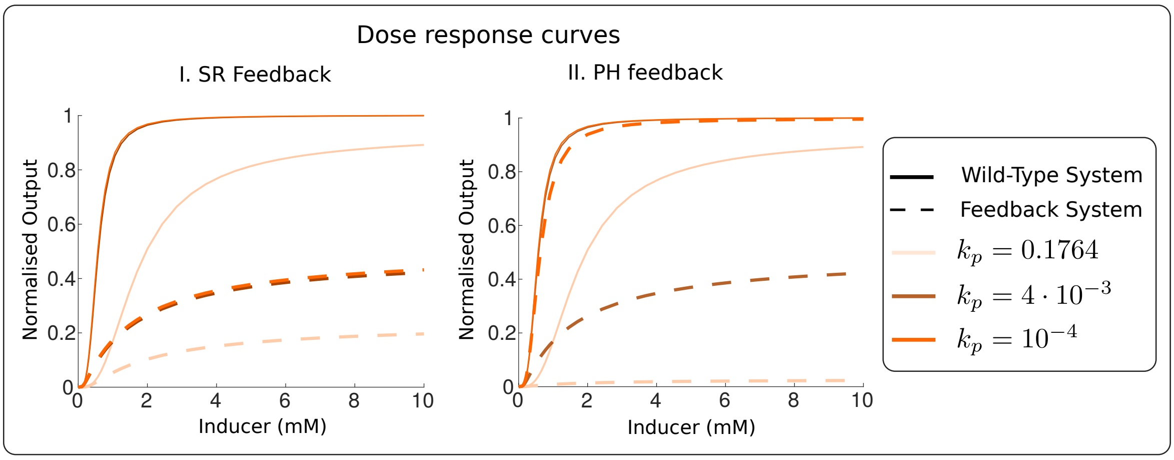

Figure S3: Dose response curves of the wild-type, the SR feedback and the PH feedback systems. In both figures the parameters are from Table S1 except for $k_p$. The SR feedback is less sensitive to changes to $k_p$, while the PH feedback exhibited similar dose response curves to the SR feedback for $k_p = 4 \cdot 10^{-3}$ $[1/(\mu\mathrm{M}\,\mathrm{min})]$, an almost unaffected inducer response for $k_p = 10^{-4}$ $[1/(\mu\mathrm{M}\,\mathrm{min})]$ and an almost complete response repression for $k_p = 1.764 \cdot 10^{-1}$ $[1/(\mu\mathrm{M}\,\mathrm{min})]$. Varying $k_t$ in the interval $[1,\ 6.12 \cdot 10^3]$ $[1/(\mu\mathrm{M}\,\mathrm{min})]$ did not yield considerable changes in the dose response curves.

#### D.3 Comparison of the Dichotomous Feedback Mechanisms

We compared the feedback mechanisms by reproducing the results in Figure 4 and Figure 5 for $P = 1$, various dephosphorylation rates $k_p$, while other parameters taken from Table S1 unless otherwise specified. For all the computation results in Figures S3, S4 and S5 varying the phosphorylation rate $k_t$ in the interval $[1,\ 6.12{\cdot}10^3]$ $[1/(\mu\mathrm{M}\,\mathrm{min})]$ yielded indistinguishable by the naked eye curves and therefore they were not plotted. Overall, these numerical computations suggest that the feedback mechanisms have similar properties, but for some large dephosphorylation rates $k_p$ the feedback strength of the phosphatase feedback is larger. This can be seen from our steady-state models (S36), and (S38), as well. Indeed, since the values of $\mathrm{RR}_{\mathrm{tot}}$, $k_t$, $k_{tc}$ are much larger than $\delta$, varying $k_t$ and $k_{tc}$ in $[1,\ 6.12 \cdot 10^3]$ $[1/(\mu\mathrm{M}\,\mathrm{min})]$ does not have a significant effect on $k_{tc}/(k_t\mathrm{RR}_{\mathrm{tot}} + \delta)$, on the other hand the values of $\mathrm{HK}_{\mathrm{tot}}$, $k_p$, $k_{pc}$ are comparable to $\delta$ and hence varying $k_p$, $k_{pc}$ can significantly increase or decrease the feedback strength as suggested in Figure S3. This also explains why intrinsic noise and the constant disturbance rejection properties are better for the phosphatase feedback with $k_p = 0.1764$ $[1/(\mu\mathrm{M}\,\mathrm{min})]$.

Finally, we computed the magnitude of the sensitivity function for the inducer concentrations $[I]$ equal to $[0.1, 1, 2, 5, 10]$ [mM], $k_t = [6.12 \cdot 10^3, 6.12 \cdot 10^2, 1]$ $[1/(\mu\mathrm{M}\,\mathrm{min})]$, $k_p = [0.1764, 4 \cdot 10^{-2}, 4 \cdot 10^{-3}]$ $[1/(\mu\mathrm{M}\,\mathrm{min})]$, and $P = [0.1, 0.4, 0.7, 1]$. In the SR feedback architecture, the maximum magnitude was equal to 1.2152, and the PH feedback architecture the maximum magnitude was equal to 1.2022. This suggests that both feedback architectures are robust according to standard engineering guidelines.

#### D.4 Modelling and Parameter Values for the Molecular sequestration based and Transcriptional Feedback Architectures

As discussed in the main text we consider the two feedback mechanisms depicted in Figures 8.I and 8.II:

(a) Molecular sequestration based feedback annihilating the kinase realised by the following reactions:

$$\begin{gathered} \emptyset \xrightarrow{k_x(\mathrm{RR_p})} X, \\ X + \mathrm{HK_p} \xrightarrow{k_{at}} \emptyset, X + \mathrm{HK} \xrightarrow{k_{at}} \emptyset. \end{gathered} \tag{S40}$$

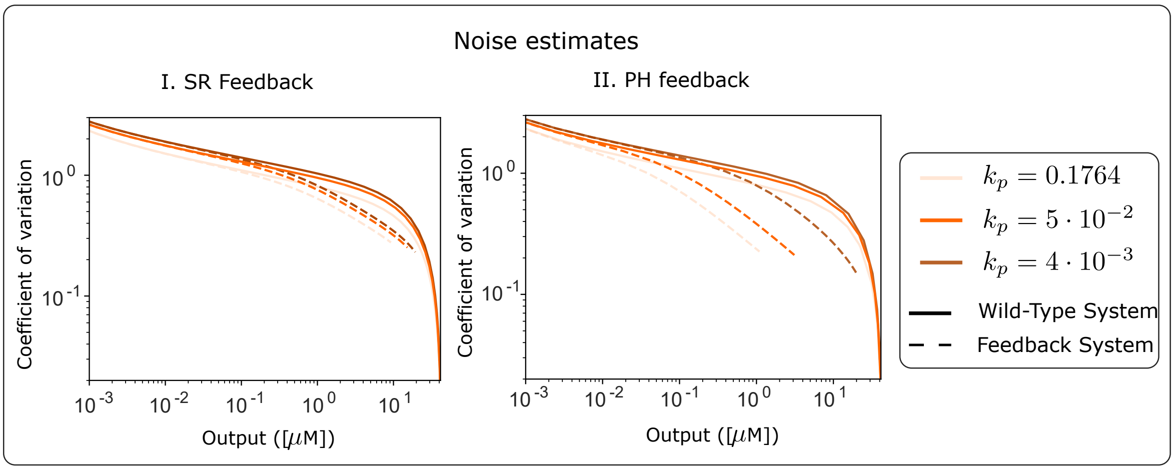

Figure S4: Coefficient of variation of the wild-type and the closed loop systems against the mean output protein concentrations. Both systems reduce the noise in a two-component system, however, the PH feedback system appears to reduce noise much better for $k_p = 0.1764$ $[1/(\mu\mathrm{M}\,\mathrm{min})]$ and $k_p = 5 \cdot 10^{-2}$ $[1/(\mu\mathrm{M}\,\mathrm{min})]$ than SR feedback systems. This is due to the higher feedback strength in the PH feedback system for these values of $k_p$. While noise attenuation for $k_p = 4 \cdot 10^{-3}$ $[1/(\mu\mathrm{M}\,\mathrm{min})]$ is similar for both feedbacks, phosphatase feedback still attenuates noise slightly better than the response regulator feedback. Varying $k_t$ in the interval $[1,\ 6.12 \cdot 10^3]$ $[1/(\mu\mathrm{M}\,\mathrm{min})]$ did not yield considerable changes in the curves.

(b) Transcriptional feedback realised by repressing the production of the histidine kinase:

$$\begin{aligned} \emptyset &\xrightarrow{k_x(\mathrm{RR_p})} X, \\ \emptyset &\xrightarrow{k_{\mathrm{HK}}(X)} \mathrm{HK}, \end{aligned} \tag{S41}$$

In both cases, $X$ is the protein realising the feedback and

$$\begin{aligned} k_{\mathrm{HK}} &= \frac{\beta_{\mathrm{HK}}}{1 + X/K_{dx}}, \\ k_x(\mathrm{RR_p}) &= P k_{\mathrm{out}}([\mathrm{RR_p}]). \end{aligned}$$

We take the parameter values from Table S1, and for the transcriptional feedback we set $K_{dx} = 6$ $[\mu\mathrm{M}]$, while for the molecular sequestration based feedback $k_{at} = 4.08 \cdot 10^{-3}$ $[1/(\mu\mathrm{M}\,\mathrm{min})]$, which is approximately equal to the dephosphorylation rate $k_p$. We selected the parameter values for the transcriptional and molecular sequestration based feedbacks so that the dose response curves would match with the SR feedback system.

Again to compute the sensitivity function, we perturb only the process outputs corresponding to $[\mathrm{RR_p}]$ and linearise the models similarly to the signal sequestration feedback systems. This gives the results depicted in Figure S6.

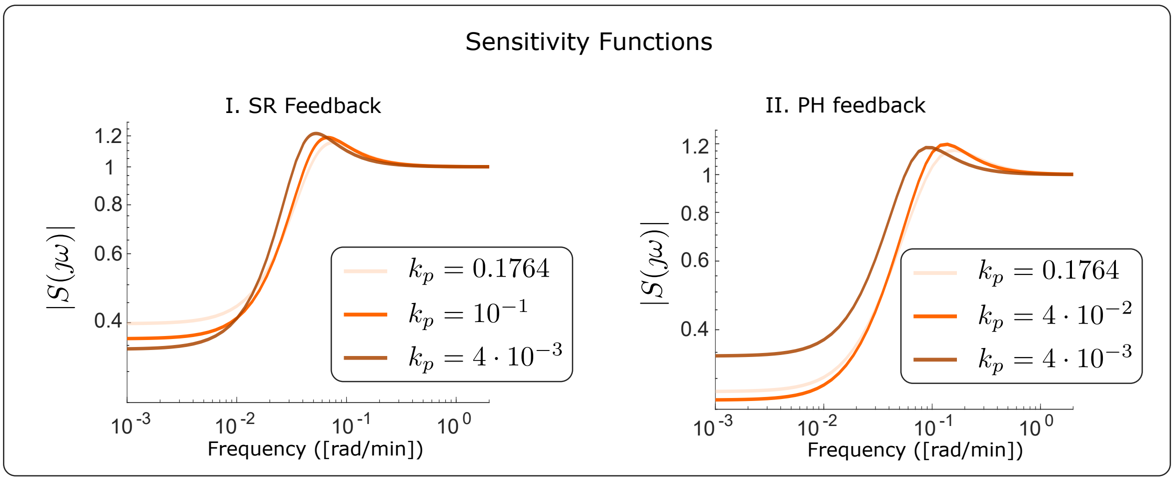

Figure S5: The magnitude of the sensitivity function of the closed loop systems for $[I] = 2$ [mM]. The shape and the values of the magnitude of the frequency response suggest that the closed loop system is robust towards perturbations. The PH feedback system appears to reject the constant disturbances better (the value $\|S(0)\|_2$ is small) than the SR feedback for $k_p = 0.1764$ $[1/(\mu\text{M min})]$ and $k_p = 4 \cdot 10^{-2}$ $[1/(\mu\text{M min})]$. This is due to the higher feedback strength in the PH feedback system for these values of $k_p$. Varying $k_t$ in the interval $[1,\ 6.12 \cdot 10^3]$ $[1/(\mu\text{M min})]$ did not yield considerable changes in the curves.

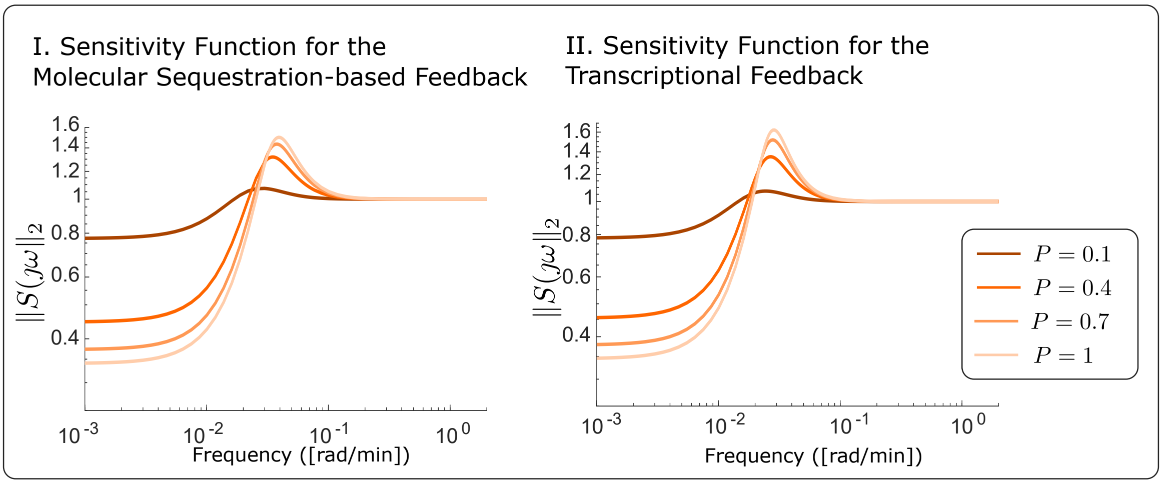

Figure S6: I. and II. Sensitivity functions for the molecular sequestration based and transcriptional feedback loops for $[I] = 2$ [mM]. The magnitude of the sensitivity functions are significantly larger than 1.2 indicating weak robustness properties.

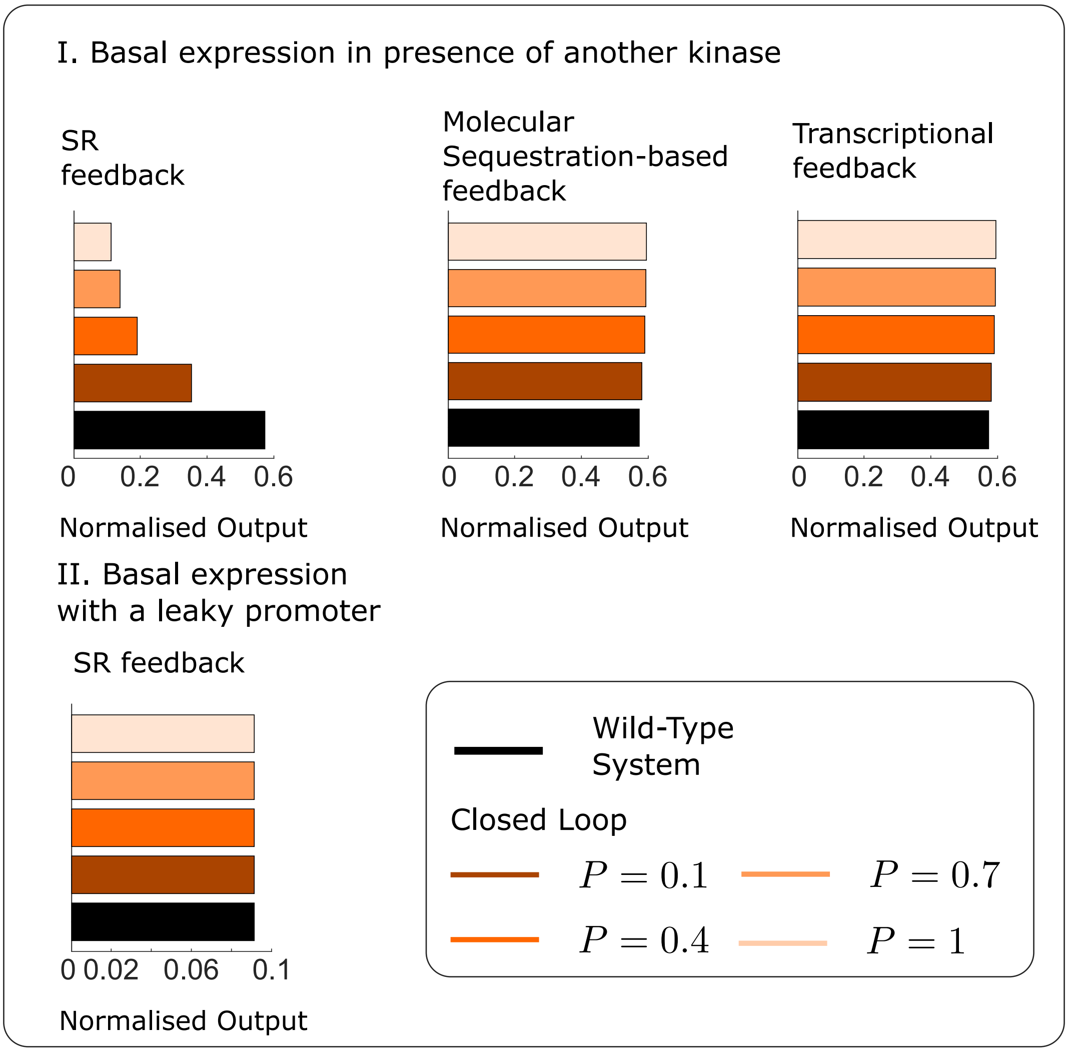

Figure S7: I. Basal expression (with $[I] = 0$ [mM]) in presence of another histidine kinase in the crosstalk system from the main text for different feedback architectures: The SR feedback system, molecular sequestration based and transcriptional feedback systems from Section D.4. The SR feedback system considerably reduces the basal expression, while molecular sequestration based and transcriptional feedback architectures have almost no effect on basal expression. II. Basal expression (with $[I] = 0$ [mM]) in presence of a leaky output promoter, i.e., the expression term for the output protein is $k_{\text{out}}([\text{RR}_\text{p}]) = 0.1 + k_{\text{out}-\max}\frac{([\text{RR}_\text{p}]/K_{dr})^n}{([\text{RR}_\text{p}]/K_{dr})^n+1}$. The SR feedback system has no effect on basal expression in presence of a leaky promoter.
